# Ultrahigh-affinity transport proteins from ubiquitous marine bacteria reveal mechanisms and global patterns of nutrient uptake

**DOI:** 10.1101/2023.02.16.528805

**Authors:** Ben E. Clifton, Uria Alcolombri, Colin J. Jackson, Paola Laurino

## Abstract

SAR11 bacteria are the most abundant members of the global ocean microbiome and have a broad impact on ocean ecosystems. To thrive in their competitive oligotrophic environments, these bacteria rely on solute-binding proteins (SBPs) that facilitate nutrient uptake through ABC transporters. Nonetheless, previous studies have been unable to access the molecular mechanisms and functions of these transporters because they rely heavily on homology-based predictions. These mechanisms and functions are essential to understand biogeochemical cycling in the ocean, including assimilation of dissolved organic matter (DOM). Here, by doing a biochemical study of the collective behavior of all SBPs in a SAR11 bacterium, we discover that these transporters have unprecedented binding affinity (*K*_d_ ≥30 pM) and unexpectedly high binding specificity, revealing molecular mechanisms for oligotrophic adaptation. Our study uncovers new carbon sources for the SAR11 bacteria and provides an accurate biogeographical map of nutrient uptake in the ocean. Our results show how functional adaptation at the molecular level in ubiquitous marine bacteria impacts global patterns of DOM assimilation and provides insight into the contribution of different compounds to oceanic nutrient cycles.

## Introduction

The sunlit surface ocean is dominated by heterotrophic bacterioplankton, particularly those belonging to the SAR11 clade of Alphaproteobacteria (*Pelagibacterales*)^1^. SAR11 bacteria are globally distributed and abundant, constituting ~20–45% of prokaryotic cells and ~18% of the biomass in the surface ocean and having an estimated global population size of 2.4 × 10^28^ cells^2,3^. Like other bacteria adapted to oligotrophic (nutrient-poor) environments, they exhibit a small size (~0.1–0.5 μm^3^), extremely streamlined genome (~1.2–1.4 Mbp), and limited metabolic versatility^1,4,5^. SAR11 bacteria rely largely on the uptake of dissolved organic matter (DOM) to meet their requirements for carbon, nitrogen, sulfur and phosphorus, and are highly active consumers of labile DOM, accounting for ~30–60% of assimilation of amino acids, taurine, glucose, and dimethylsulfoniopropionate (DMSP) in the surface ocean^6–10^. Due to their high abundance, SAR11 bacteria have major biogeochemical importance; for example, they produce climate-active gases such as methane^7^ and dimethyl sulfide^11^, contribute to carbon sequestration through the microbial carbon pump^12^, and divert carbon from the biological carbon pump through respiration of dissolved organic carbon (DOC)^13^. Thus, understanding the physiology and metabolic capabilities of SAR11 bacteria is critical to our understanding of marine ecosystems.

To compete for nutrients in the oligotrophic ocean environment, SAR11 bacteria rely heavily on ATP-binding cassette (ABC) transporters to facilitate nutrient uptake. These transporters represent the most abundant superfamily of transport proteins used for high-affinity nutrient uptake in Gram-negative bacteria, and are composed of a periplasmic SBP, two transmembrane domains, and two cytoplasmic nucleotide-binding domains (**Fig. 1**)^14^. SBPs bind their substrates with high affinity in the periplasm (*K*_d_ typically in the nM–μM range^15^) and deliver them to the transmembrane complex of the ABC transporter, which couples binding and hydrolysis of ATP to translocation of the substrate across the inner membrane. Consistent with the physiological importance of ABC transporters for high-affinity nutrient uptake, SAR11 bacteria devote a large proportion of their streamlined proteome to ABC transporters^16,17^; for example, SBPs represented ~67% of SAR11-derived spectra in metaproteomic analysis of environmental samples from the Sargasso Sea^17^.

**Fig. 1.**
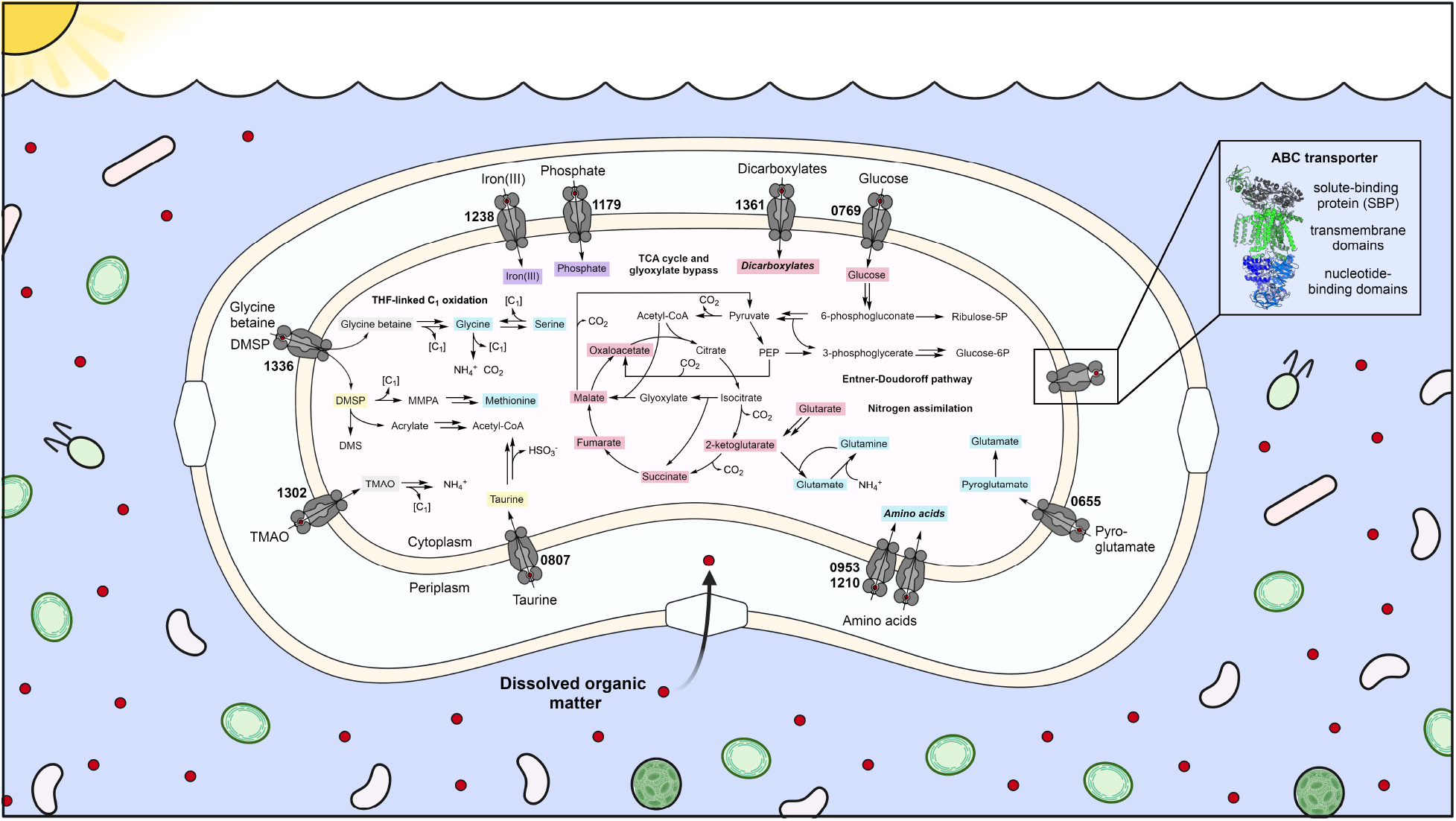
Ubiquitous SAR11 bacteria use ABC transporters to uptake dissolved organic matter from the ocean, playing a critical role in oceanic cycles. The enlarged SAR11 bacterium shows the ABC transporters of ‘*Ca*. P. ubique’ HTCC1062 and their relationship with key pathways of carbon, nitrogen, and sulfur metabolism. Metabolites identified as SBP ligands in this work are highlighted in color: blue, amino acids and their derivatives; red, carbon sources; yellow, sulfur sources; grey, C_1_ sources; purple, other. Metabolic pathways are based on reactions proposed in this work (**Fig. S11**) and refs ^27,28,38,40,82,83^. [C_1_] refers to a C_1_ unit oxidized to CO_2_ *via* the THF-linked oxidation pathway. Abbreviations: DMS, dimethylsulfide; DMSP, dimethylsulfoniopropionate; glucose-6P, glucose-6-phosphate; MMPA, methylmercaptopropionate; PEP, phosphoenolpyruvate; ribulose-5P, ribulose-5-phosphate; THF, tetrahydrofolate; TMAO, trimethylamine-*N*-oxide.

The high abundance and nutrient uptake activity of SAR11 bacteria in the ocean, combined with the abundance of ABC transporters in these bacteria, suggests that a small number of transport proteins in SAR11 bacteria make an outsized contribution to global assimilation of labile DOM in the ocean. However, the properties of these transporters and their specific functions (i.e., transported metabolites) are mostly unknown, limiting our knowledge of the full range of DOM that can be assimilated by SAR11 bacteria, nutrient exchange within the marine microbial community, and the molecular mechanisms for high-affinity nutrient uptake. Although homology-based predictions are available, homology-based predictions of protein function have limited accuracy, especially for functionally diverse protein superfamilies like ABC transporters^18,19^. Transport proteins can also be characterized experimentally through radioassays of nutrient uptake in cultured cells, and this approach has been used to characterize the broad-specificity osmolyte transporter of the SAR11 bacterium ‘*Candidatus* Pelagibacter ubique’^20^. However, the difficulty of cultivating slow-growing and fastidious SAR11 bacteria makes this a challenging approach for high-throughput characterization of ABC transporters. Furthermore, due to the genetic intractability of SAR11 bacteria, the observed transport activity cannot be linked to specific ABC transporter genes, limiting integration of the resulting physiological data with existing multi-omics datasets to uncover the broader geochemical and ecological significance of transport activity. An alternative approach to elucidate the functions of ABC transporters is through biochemical characterization of the corresponding SBPs^21^, which relies on the fact that the specificity and affinity of nutrient uptake by ABC transporters is mainly determined by the binding specificity and affinity of the corresponding SBPs^22^, and has been demonstrated to be a valuable method for discovery of new metabolic pathways, particularly by the Enzyme Function Initiative project^23,24^.

Here, we used this approach to systematically interrogate the function of all ABC transporters in the genome of the prototypical SAR11 bacterium ‘*Ca*. P. ubique’ strain HTCC1062. Using high-throughput screening together with rigorous structural and biophysical characterization, we identified the function of the majority of these ABC transporters. Revision of incorrect homology-based functional predictions allowed us to obtain accurate biogeographical maps of nutrient uptake and identify new transport capabilities and potential carbon sources for SAR11 bacteria. In particular, we identified a high-affinity, broad-specificity transporter for C_4_/C_5_ dicarboxylates that is widely found among SAR11 ecotypes and abundantly distributed in metagenomic and metatranscriptomic datasets, implicating these dicarboxylates as major physiologically relevant carbon sources. Finally, we show how the identification of systematic trends in SBP properties, including their extremely high binding affinity, unexpectedly high binding specificity, and limited functional redundancy, provides insight into the evolutionary success of SAR11 bacteria in the oligotrophic ocean environment.

## Results and Discussion

Fourteen ABC SBPs were identified through genomic analysis of ‘*Ca*. P. ubique’ strain HTCC1062 (**Materials and Methods**, **Table S1**); these SBPs are found widely across SAR11 bacteria, and conversely, the most abundant ABC SBPs across SAR11 bacteria are represented in this strain (**Fig. S1**). Expression of most of these SBPs in cultured and/or environmental SAR11 cells has been previously demonstrated by proteomic analysis (**Table S2**)^16,17^. Twelve of the SBPs yielded soluble protein upon heterologous expression in *Escherichia coli* strains BL21(DE3) or SHuffle T7 and were successfully purified, while the remaining two proteins, SAR11_0271 and SAR11_1346, could not be expressed in soluble form under any tested condition, nor refolded *in vitro* from insoluble material (**Supplementary Methods**). Two of the SBPs, SAR11_1179 and SAR11_1238, were predicted to represent proteins that are found widely in bacteria and bind inorganic solutes with high specificity: phosphate and iron(III), respectively. Thus, the functional predictions for these two proteins were directly tested and confirmed by differential scanning fluorimetry (DSF) and isothermal titration calorimetry (ITC) (SAR11_1179; **Fig. S2**) or UV-vis spectroscopy (SAR11_1238, **Fig. S3**) rather than high-throughput screening.

In the remaining cases, the function of each SBP was first identified by high-throughput screening of metabolite libraries by DSF and further validated by ITC. Firstly, the target protein was screened by DSF against a commercially available metabolite library, representing a set of ~330 unique metabolites, including many common carbon, nitrogen, phosphorus, and sulfur sources (see **Table S3** for full list). This library was supplemented with a manually curated set of ~30 metabolites that are known to be important for ‘*Ca*. P. ubique’ and other marine bacteria (e.g., osmolytes, sulfonates, and vitamin derivatives)^25–28^ or that were considered to be potential ligands based on the computational annotations of SBP function (e.g., opines). Metabolites that resulted in a *T*_M_ increase (Δ*T*_M_) of ≥2 °C by DSF were considered to be potential ligands (**Table S4**)^29^. Secondly, a representative subset of the resulting hits was selected, and binding of this subset of ligands to the target protein was confirmed and rank-ordered by repeating DSF with each ligand at a fixed concentration (10 mM) (**Fig. 2**). Finally, to provide further evidence that the observed increases in *T*_M_ were a result of specific, high-affinity protein-ligand interactions rather than non-specific protein stabilization, the DSF experiments were repeated with a range of ligand concentrations (**Fig. S4**). Using this workflow, putative functions were identified for eleven SBPs, i.e., all proteins that could be expressed and purified, except SAR11_1068. We showed previously that SAR11_1068 does not have the annotated function (cyclohexadienyl dehydratase activity) and reported extensive but ultimately unsuccessful efforts to identify its function^30^. The protein was subjected to further high-throughput screening in this work, but no potential ligands were identified.

**Fig. 2.**
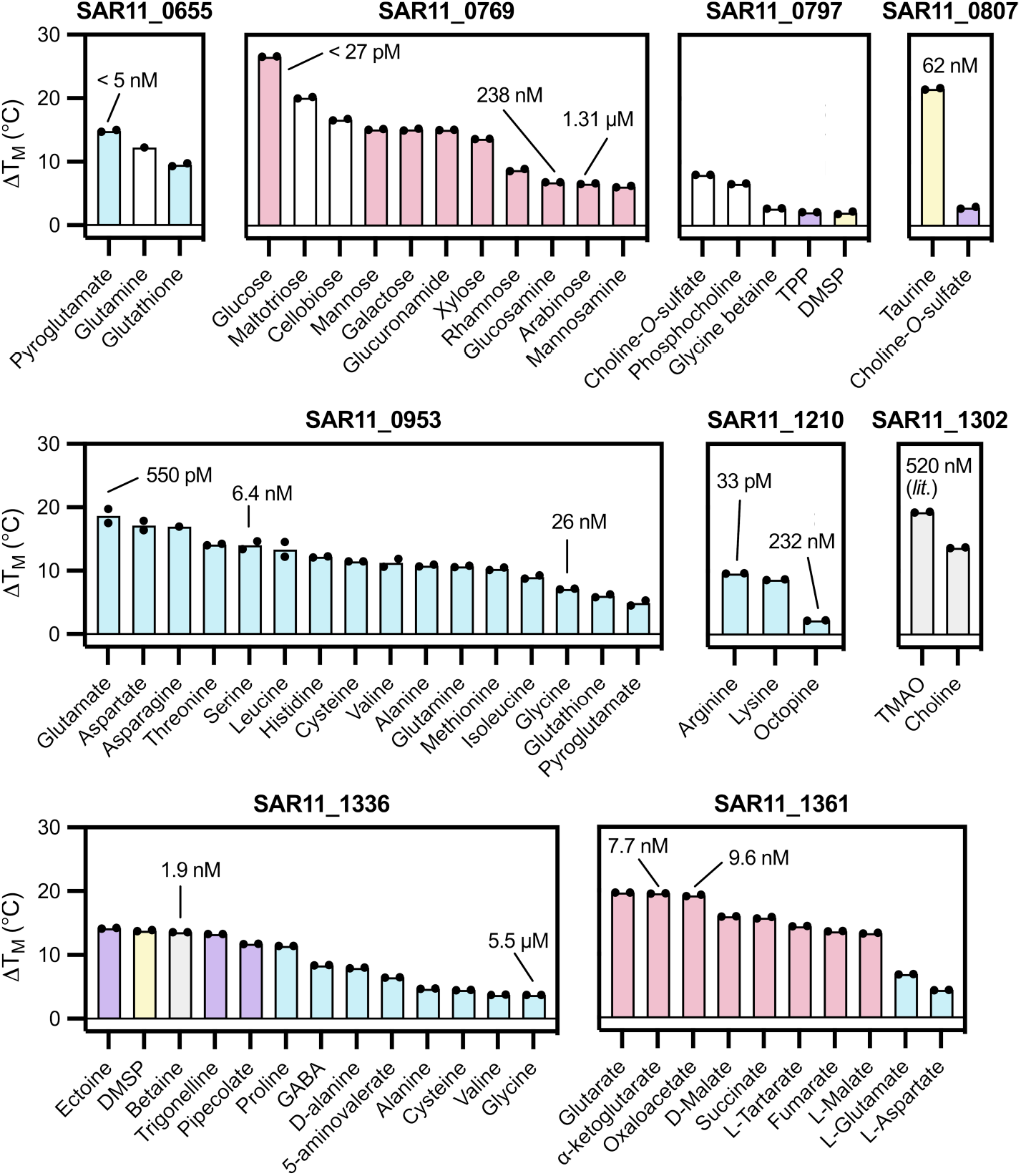
SBPs from SAR11 bacteria (’ *Ca*. P. ubique’ HTCC1062) show unexpectedly high binding specificity. We measured the change in denaturation temperature (Δ*T*_M_) in the presence of 10 mM ligand by DSF. Ligands that resulted in a significant increase in *T*_M_ (≥ 2°C) at a concentration of 10 mM are shown. Protein-ligand interactions that were verified by ITC are labelled with the measured *K*_d_ value for the interaction. Ligands are colored according to **Fig. 1**; white columns indicate ligands that showed a significant increase in *T*_M_ in DSF but did not show binding in ITC. Columns represent the mean of two technical replicates shown as individual data points. The *K*_d_ value for SAR11_1302 was taken from ^31^. Abbreviations: DMSP, dimethylsulfoniopropionate; TMAO, trimethylamine-*N*-oxide; TPP, thiamine pyrophosphate; GABA, γ-aminobutyrate.

Next, the function of each SBP was confirmed by ITC, enabling accurate quantification of binding affinity, which is important to establish the physiological relevance of the observed protein-ligand interactions. To enable approximate correlation of the Δ*T*_M_ values obtained by DSF with binding affinity, we typically performed titrations for at least two ligands for each protein, for a total of 24 protein-ligand interactions. Using ITC, most of the protein-ligand interactions identified by DSF could be verified (**Fig. 2**, **Fig. S5**, **Table S5**), and at least one high-affinity ligand (*K*_d_ <200 nM) could be identified for each SBP, except SAR11_0797 (**Supplementary Note 1**). The interaction between SAR11_1302 and trimethylamine-*N*-oxide (TMAO) was confirmed by ITC in another report while this work was in progress^31^. Thus, in

total, ten of the fourteen SBPs of ‘*Ca*. P. ubique’ HTCC1062 were able to be confidently assigned a binding function (**Fig. 1, Table S6**). The physiological relevance of these binding functions is highlighted by the identification of SBPs that bind known nutrients of ‘*Ca*. P. ubique’ HTCC1062 such as amino acids, D-glucose, DMSP, and taurine with high affinity^6–10^.

The SBPs of ‘*Ca*. P. ubique’ HTCC1062 showed remarkably high binding affinity, with *K*_d_ ranging from 133 nM to ~30 pM (**Fig. 3A-B**). Several SBPs showed *K*_d_ values of <5 nM, which are below the quantitation limit of direct ITC experiments^32^; thus, to obtain accurate *K*_d_ values for these interactions, we also performed competitive ITC binding experiments (**Table S5**). In the case of SAR11_1210, titration with L-arginine in the presence of D-octopine as a competing ligand indicated a *K*_d_ value between 10 pM and 100 pM for L-arginine, but variable results were obtained with different concentrations of D-octopine (**Data S1**). Thus, to confirm the high affinity of this interaction, we also performed a protein-protein competition experiment in which SAR11_1210 was mixed with a previously characterized arginine-binding protein, ArgT from *Salmonella enterica* (*K*_d_ 15 nM), and then titrated with L-arginine. Fitting the resulting data to a two-sets-of-sites binding model yielded a *K*_d_ of 33 pM for the interaction between SAR11_1210 and L-arginine (**Fig. 3C**). We also solved the crystal structure of SAR11_1210 complexed with L-arginine, which showed an unusual binding mode involving a direct interaction between the ligand and the flexible hinge region linking the two α/β domains of the SBP, suggesting a possible structural basis for the high binding affinity (**Fig. 3E, Fig. S6, Supplementary Note 2**). Finally, in the case of SAR11_0769, titration with D-glucose reproducibly yielded a biphasic binding isotherm, which most likely reflects differential binding the α and β anomers of D-glucose (**Fig. 3D**), as supported by a crystal structure of SAR11_0769 complexed with β-D-glucose (**Supplementary Note 3, Fig. 3F**). Fitting the ITC data to a competitive binding model allowed estimation of the upper limit of *K*_d_ (lower limit of affinity) for the high-affinity anomer as ~27 pM. A systematic survey of literature data (*n* = 192 SBPs) revealed that the typical range of ABC SBP *K*_d_ values for organic solutes is ~10–1000 nM, with a lower limit of ~200–400 pM (mean ± s.d. of log_10_ *K*_d_ values −6.81 ± 1.13, **Fig. 3A**). Altogether, these results provide robust evidence that some SBPs in ‘*Ca*. P. ubique’ HTCC1062 exceed the previously established limits of SBP affinity for organic solutes.

**Fig. 3.**
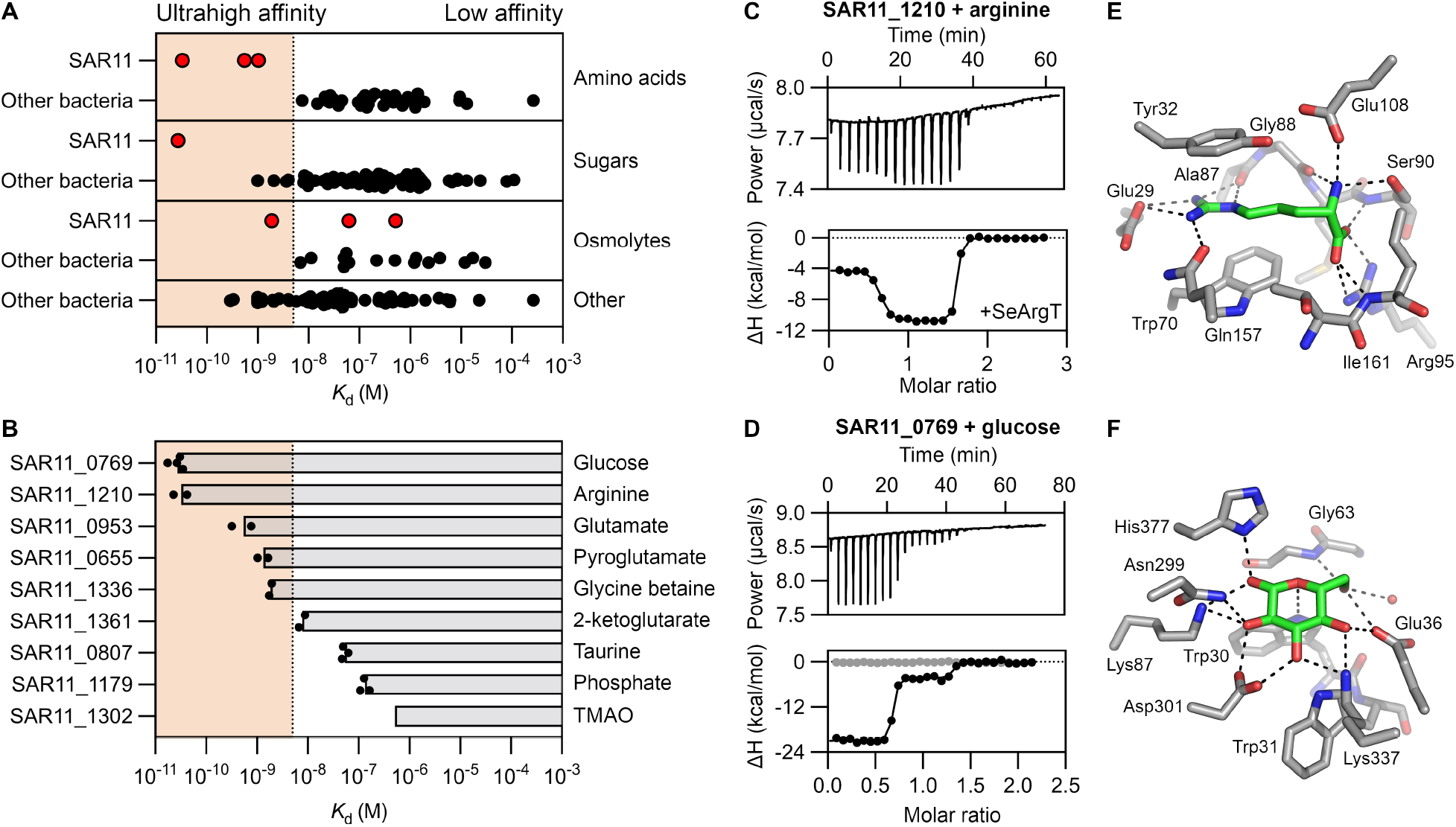
SBPs from SAR11 bacteria (’ *Ca*. P. ubique’ HTCC1062) exhibit ultrahigh binding affinity. (**A**) We compare the *K*_d_ values for SBPs from ‘*Ca*. P. ubique’ HTCC1062 with bacterial ABC SBPs previously reported in the literature (*n* = 192, **Data S2**). In cases where the SBP binds multiple ligands, data for the highest affinity ligand is shown. SBPs with a *K*_d_ <5 nM are highlighted by red shading. (**B**) We report the *K*_d_ values for highest affinity interaction for each SBP from ‘*Ca*. P. ubique’ HTCC1062. Values for SAR11_0655 and SAR11_0769 represent upper bounds on *K*_d_ (lower bounds on affinity). Bars represent mean of 2–4 technical replicates (independent titrations), shown as individual data points. (**C**-**D**) Determination of binding parameters for high-affinity interactions through competitive ITC experiments. For graphical purposes, data fitting was performed in MicroCal PEAQ software. (**C**) Simultaneous titration of SAR11_1210 and SeArgT with L-arginine. (**D**) Titration of SAR11_0769 with D-glucose, showing a biphasic binding curve. The negative control titration (buffer + D-glucose) is also shown in grey. (**E**) Binding mode of L-arginine in crystal structure of SAR11_1210 (1.32 Å). (**F**) Binding mode of D-glucose in crystal structure of SAR11_0769 (1.86 Å).

### Re-assignment of ABC transporter function enables accurate multi-omics analysis

Comparison of the experimentally determined functions of each SBP with the homology-based predictions indicated that the accuracy of the predictions was limited (**Table S6**). The binding specificities of three proteins, SAR11_0807, SAR11_1179, and SAR11_1238, were correctly predicted as taurine, phosphate, and iron(III), respectively. The predictions of SAR11_0769 and SAR11_0953 as a sugar-binding protein and general amino acid-binding protein were broadly correct, although experimental characterization enabled identification of specific ligands. On the other hand, seven of the twelve testable functional annotations were incorrect; in five of these cases, the binding specificity could be determined experimentally. For example, SAR11_1336 (*potD*), which was annotated as a spermidine/putrescine-binding protein, showed broad specificity for glycine betaine, DMSP, and other osmolytes. The binding specificity of this protein matches the transport activity of a broad-specificity osmolyte transporter previously characterized *in vivo*, which was putatively attributed to SAR11_0797 (*proX*)^20^. Overall, these results show that the ABC transporters of SAR11 bacteria transport a narrower range of nitrogen sources and broader range of carbon sources and exhibit less functional redundancy than predicted^12^, and provide further evidence that computational predictions of ABC transporter function must be treated with caution.

Importantly, the assignment of transport capabilities to specific genes enables integration of the functional data with existing genomic, transcriptomic, and proteomic data. For example, to evaluate the geographical distribution of various transport capabilities across SAR11 bacteria, we performed biogeographical analysis using metagenome and metatranscriptome datasets from the *Tara* Oceans project^33^. Consistent with the global abundance of SAR11 bacteria and the broad distribution of the characterized SBPs in SAR11 bacteria (**Fig. S1**), most of the SBP genes were present at high abundance (≥1% of total mapped reads in some cases) in the metagenome and metatranscriptome datasets, including surface, deep chlorophyll maximum (DCM) and mesopelagic samples (**Fig. 4A-B, Fig. S7-S9**). Transporters for taurine, amino acids, TMAO, glycine betaine, DMSP, and dicarboxylates showed a near-universal distribution and particularly high abundance, whereas transporters for L-pyroglutamate, phosphate, iron(III) and D-glucose showed a geographically limited distribution (**Fig. S7-S8**). These results are consistent with the known contribution of SAR11 bacteria to uptake of taurine, amino acids, and DMSP across different environments, compared with ecotype-specific and geographically variable uptake of D-glucose^1^. Similar patterns of SBP gene abundance were typically observed in the metagenome and metatranscriptome datasets, consistent with high constitutive expression and limited transcriptional regulation of most SBP genes in SAR11 bacteria, with the exception of SBPs for phosphate and iron, which showed higher expression in regions of known phosphate and iron limitation^34,35^. Notably, these interpretations of the metagenomic and metatranscriptomic data are contingent on accurate functional annotation; for example, misidentification of the osmolyte transporter as SAR11_0797 would suggest a much more limited role for DMSP and glycine betaine uptake and broader role for polyamine uptake across marine bacteria^36^.

**Fig. 4.**
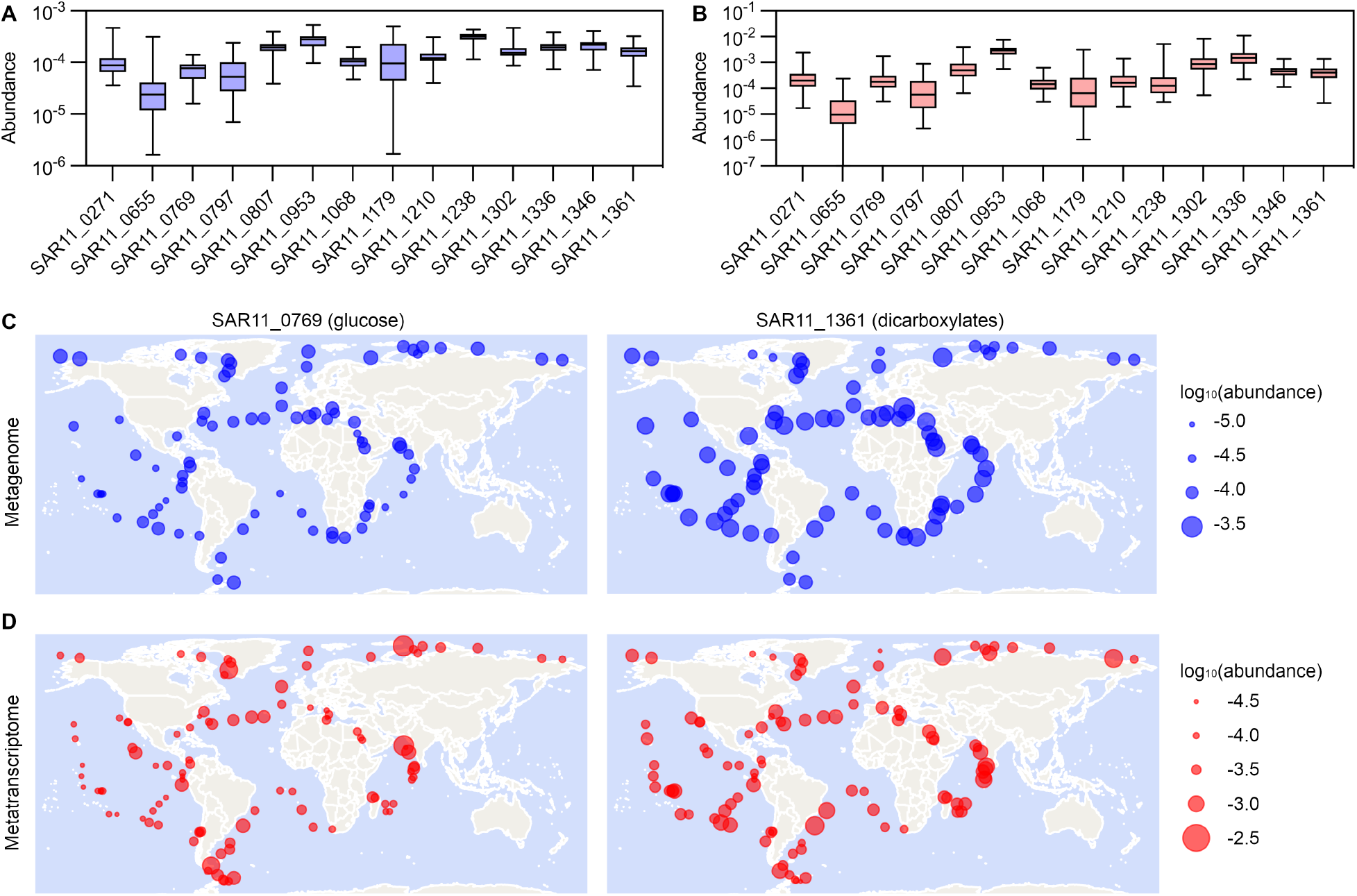
SBPs from SAR11 bacteria are abundantly distributed across the surface ocean. Abundance of SBP genes from ‘*Ca*. P. ubique’ HTCC1062 in metagenomic (**A, C**) and metatranscriptomic (**B, D**) datasets across surface samples from the *Tara* Oceans project. Data were obtained using the Ocean Gene Atlas v2.0 with an e-value cut-off of 10^−40^. Abundance is expressed as the percentage of mapped reads in each sample and is represented by point area on a linear scale. (**A**-**B**) Summary box-and-whisker plot of abundance data for each gene across surface samples. (**C**-**D**) Geographical distribution of SAR11_0769 and SAR11_1361.

### Identification of new transport capabilities for pyroglutamate and dicarboxylates

In addition to identifying transporters for known nutrients, functional characterization of SBPs also allowed identification of new transport capabilities. SAR11_0655 (L-pyroglutamate) and SAR11_1361 (C_4_ and C_5_ dicarboxylates) represent new classes of ABC transporters and previously unknown transport capabilities of SAR11 bacteria. L-Pyroglutamate, which binds to SAR11_0655 with *K*_d_ <5 nM, was a surprising ligand, as a non-proteogenic amino acid not known to be a significant component of DOC^13^. Although the occurrence of SAR11_0655 is limited among SAR11 ecotypes and mainly restricted to high latitudes (**Fig. S1**, **Fig. S7, Fig. S10**), other SAR11 bacteria appear to achieve L-pyroglutamate uptake using an alternative transporter (**Supplementary Note 4**), suggesting that L-pyroglutamate is widely utilized. Analysis of genome context suggested a pathway for utilization of exogenous L-pyroglutamate as a source of L-glutamate (**Fig. S11**). The fact that SAR11 bacteria, with their extremely streamlined genome, retain specific and high-affinity transporters for L-pyroglutamate indicates that this amino acid must be a widely available and useful carbon/nitrogen source in the ocean. More generally, given the significant challenges of identifying environmentally important metabolites in heterogeneous, dilute, and variable DOC^13^, this result suggests that identification of new transport capabilities from characterization of SBPs from oligotrophic marine bacteria might be a useful approach to identify new environmentally significant ocean metabolites from the DOC pool.

SAR11_1361 showed binding of a broad range of dicarboxylates that participate in the tricarboxylic acid (TCA) cycle (**Fig. 1, Fig. 2**). This gene is known to be associated with carbon starvation in SAR11 bacteria; transcription and/or expression is upregulated upon carbon limitation in the dark (i.e., energy-starved conditions)^37^ and downregulated upon nitrogen and sulfur limitation^38,39^. Analysis of genome context also suggested a potential pathway for utilization of exogenous glutarate (**Fig. S11**), which was subsequently confirmed as a ligand of SAR11_1361 by DSF (**Fig. 2**). These results suggest a broad capacity of ‘*Ca*. P. ubique’ HTCC1062 to assimilate dicarboxylates. Genomic and biogeographical analysis indicated that this capability is also widely distributed among SAR11 bacteria: the dicarboxylate transport protein SAR11_1361 shows a broader distribution among published SAR11 genomes than the glucose transport protein SAR11_0769 (**Fig. S1**), and shows a broader geographical distribution in the *Tara* Oceans metagenome and metatranscriptome datasets, including both coastal and open ocean samples (**Fig. 4C, D**), despite the fact that SAR11_1361 has a much more limited phylogenetic distribution among bacteria (**Fig. S12**). In the context of uncertainty surrounding the carbon sources that are universal to SAR11 bacteria^40^ (**Supplementary Note 5**), the identification of an SBP with high affinity (*K*_d_ <10 nM) and broad specificity for C_4_/C_5_ dicarboxylates that is conserved among SAR11 ecotypes despite stringent genome streamlining, and widely distributed and highly transcribed throughout the ocean, provides strong evidence that these dicarboxylates are physiologically important carbon sources in SAR11 bacteria.

### The SBPs of ‘ *Ca*. P. ubique’ show mixed specificity and ultrahigh affinity

Systematic characterization of SBPs provided a global view of transporter specificity and affinity in ‘*Ca*. P. ubique’ HTCC1062, providing insight into the physiology of oligotrophic bacteria. It has long been hypothesized that oligotrophic bacteria with streamlined genomes, including SAR11 bacteria, rely on broad-specificity transporters to enable transport of a broad range of nutrients with a limited number of transporters^4^. Indeed, a broad-specificity osmolyte transporter was already identified^20^, and two more broad-specificity SBPs for amino acids and dicarboxylates were characterized in this work. However, the majority of SBPs (7/10) showed high binding specificity, suggesting a more nuanced view of uptake specificity in oligotrophic bacteria. Genome streamlining results in reduction of metabolic genes in addition to transporter genes; thus, broad-specificity ABC transporters are associated with a risk of futile uptake of metabolites that cannot be utilized, especially given the high compositional complexity of ocean DOC. Our results show that ‘*Ca*. P. ubique’ HTCC1062 is highly selective in its nutrient uptake, using a small number of broad-specificity transporters mainly for metabolites that can be utilized without dedicated catabolic pathways, including amino acids and TCA cycle intermediates. The remaining transporters show high specificity and mainly cover specific gaps in the broad-specificity transporters; indeed, there is very little redundancy in binding specificity between the SBPs. The high specificity of these transporters does not result from a negative tradeoff between specificity and affinity; for example, the broad-specificity amino acid-binding protein is estimated to have nanomolar affinity for ~15 proteinogenic amino acids (based on measured Δ*T*_M_ and *K*_d_ values), with a maximum of 550 pM for L-glutamate, demonstrating that broad specificity is compatible with high affinity. Furthermore, two of the three broad-specificity transporters (SAR11_0953 and SAR11_1336) are widely distributed among Proteobacteria, indicating that use of broad-specificity transporters is not unique to oligotrophic bacteria (**Fig. S10**). Overall, these considerations suggest that oligotrophic bacteria likely show greater selectivity in nutrient uptake than previously assumed.

Our results revealed that a systematic increase in SBP binding affinity is a major adaptation of ‘*Ca*. P. ubique’ HTCC1062 to low nutrient concentrations in the oligotrophic environment. The binding affinity of the ‘*Ca*. P. ubique’ HTCC1062 SBPs was remarkably high on average, and substantially exceeded the known range of SBP binding affinity in some cases (**Fig. 3A**). *K*_d_ values in the picomolar-low nanomolar range were observed in most cases, in concordance with the picomolar-low nanomolar concentrations of amino acids and other nutrients typically observed in the surface oligotrophic ocean (see observed concentrations in **Table S7**; ^41–43^) and picomolar-low nanomolar uptake affinities (specifically, *K*_s_ + [S] values) for various metabolites in environmental samples from the surface ocean^41,44^, for which the corresponding transport proteins have generally not been identified^20^. In addition to the known correlation between SBP binding affinity and ABC transporter uptake affinity^22^, the physiological relevance of the observed binding affinity in SBPs is indicated by several considerations: (1) the observed *K*_d_ of 1.9 nM for the interaction between SAR11_1336 and glycine betaine is in excellent agreement with the previously measured *K*_s_ value for the corresponding transporter (0.89 nM; ^20^); (2) mathematical models of ABC transporter activity indicate that uptake affinity should be greater than SBP binding affinity when SBP concentration is high (as in SAR11 bacteria)^45^; and (3) the binding affinity of an SBP has physiological significance itself, because it determines the concentration at which nutrients can be accumulated in the periplasm^46^. Whereas previous work has shown how extreme selective pressure driven by large population size under low-nutrient conditions has driven systematic adaptation of SAR11 bacteria at the genome and cellular levels (for example, reduction of GC content^4^ and increase in periplasmic volume^47^), this work shows that systematic adaptation of the biophysical properties of SBPs is another important factor in the evolutionary success of SAR11 bacteria.

The identification of SBPs with unprecedented binding affinity in the genome of ‘*Ca*. P. ubique’ HTCC1062 resolves uncertainty about the discrepancy between the observed affinity of nutrient uptake by microbial communities in the ocean and the binding affinity of previously characterized nutrient uptake transporters. To explain this apparent discrepancy, various alternative mechanisms for high-affinity nutrient uptake in oligotrophic bacteria have been proposed. For example, a recent modelling study showed that the uptake affinity of an ABC transporter depends on both SBP concentration and binding affinity, and suggested that oligotrophic bacteria might use high SBP expression to achieve high uptake affinity without increasing SBP binding affinity^45^; in contrast, our results show that high uptake affinity can be explained without accounting for periplasmic SBP concentration. As another example, the observation that the binding affinity of known phosphate-binding proteins (~1 μM) is much higher than concentrations of inorganic phosphate in phosphate-depleted regions (<5 nM) led to the proposal of an alternative mechanism for accumulation of inorganic phosphate in the periplasm of oligotrophic bacteria^46,48^. Interestingly, the phosphate-binding protein of ‘*Ca*. P. ubique’ HTCC1062 does indeed have relatively low binding affinity (133 nM), which may reflect the challenge of discriminating phosphate from sulfate, which is present at a concentration of ~28 mM in the ocean; although phosphate-binding proteins from sulfate-rich environments can achieve a discrimination factor of >10^5^ (ref. 49), there is presumably a biophysical limit on discrimination of these two anions due to their physicochemical similarity. Consistent with the hypothesis that the binding affinity of phosphate-binding proteins is constrained by the requirement for discrimination of phosphate and sulfate, SAR11_1179 showed a small but significant decrease in apparent binding affinity in the presence of 28 mM sulfate (6.7-fold decrease to 890 nM, *P* < 0.0001, two-tailed *t*-test on log_10_ *K*_d_ values, **Fig. S2**).

## Conclusion

Systems-level approaches based on metatranscriptomics and related methods are highly valuable for profiling of biological function in complex microbial communities across different environments, providing insight into their ecological and biogeochemical functions^50–52^. However, a limitation of these methods is that they depend on homology-based predictions of protein function, which vary dramatically in accuracy between protein families and are usually not validated^18^. Here, we have shown how targeted functional characterization of environmentally abundant proteins can be integrated with existing multi-omics and physiological data to provide insight over multiple biological scales, ranging from mechanisms of functional adaptation at the molecular level to global patterns of ocean nutrient uptake. We anticipate that improved computational annotation and continued experimental annotation of protein function will be essential to extract maximal value from increasingly high-resolution ocean microbiome datasets and fulfil the broader goal in microbial ecology of bridging microbial gene function and ocean ecosystems biology on a planetary scale^35,53^.

## Materials and Methods

### Identification of SBP genes

Fifteen candidate SBP genes in the genome of ‘*Candidatus* Pelagibacter ubique’ strain HTCC1062 were identified through a search of the TransportDB 2.0 database (http://membranetransport.org; accessed 22 January 2020)^54^. One of these genes, SAR11_0371, was annotated as a “possible transmembrane receptor” in UniProt and showed a non-canonical predicted domain structure consisting of a short SBP-like domain (170 amino acids) followed by a coiled coil domain and unidentified C-terminal domain. Additionally, genome context analysis showed that, unlike the other SBP genes in ‘*Ca*. P. ubique’, SAR11_0371 was not colocalized with genes encoding the membrane permease or ATP-binding cassette components of an ABC transport system. Thus, SAR11_0371 was considered not to represent an SBP component of an ABC transport system and was excluded from the analysis. We also attempted to identify additional SBP genes through a search of the UniProt database for proteins in ‘*Ca*. P. ubique’ belonging to Pfam clans CL0177 (PBP; periplasmic binding protein) and CL0144 (Periplas_BP; periplasmic binding protein like); however, this search did not return any additional candidate genes.

### Cloning

The protein sequence of each SBP was obtained from the UniProt database. Signal sequences were predicted using the SignalP 5.0 server^55^ and removed. The protein sequences were then back-translated and codon-optimized for expression in *Escherichia coli*, and the resulting genes were obtained as synthetic DNA from Twist Bioscience (South San Francisco, CA) or Integrated DNA Technologies (Coralville, IA). The synthetic genes were cloned into the NdeI/XhoI site of the pET-28a(+) expression vector by In-Fusion cloning using the In-Fusion HD Cloning Kit (Takara Bio, Kusatsu, Japan), yielding expression constructs with an N-terminal hexahistidine tag and thrombin tag. Correct assembly of each expression vector was confirmed by Sanger sequencing (FASMAC, Atsugi, Japan). The sequences of oligonucleotides and synthetic genes used in this study are listed in **Table S8**.

### Optimization of protein expression

Protein expression was initially tested in *E. coli* BL21(DE3) cells grown in Luria-Bertani (LB) and Terrific Broth (TB) media at 30 °C and 17 °C. SAR11_0655 showed optimal soluble expression in LB medium at 17 °C, while seven proteins (SAR11_0797, SAR11_0807, SAR11_1068, SAR11_1179, SAR11_1210, SAR11_1238, and SAR11_1361) showed optimal soluble expression in TB medium at 17 °C. The remaining proteins, which showed low or nil soluble expression under these conditions, were tested for expression in *E. coli* BL21(DE3) cells grown in LBNB (LB + 0.2% (w/v) glucose, 0.5 M NaCl, and 1 mM betaine) and LBE (LB + 3% (v/v) ethanol) media, which have been reported to induce chaperone expression and promote soluble protein expression^56,57^; however, significant improvements in expression were not observed. Finally, proteins were tested for expression in *E. coli* SHuffle T7 cells (New England Biolabs, Ipswich, MA) in TB medium at 17 °C; this strain expresses the disulfide bond isomerase DsbC, which can increase soluble recombinant expression of cytoplasmic proteins by promoting correct formation of disulfide bonds. Soluble expression of SAR11_0769, SAR11_0953, SAR11_1302, and SAR11_1336 was achieved under these conditions. Protein expression under each condition was evaluated by SDS-PAGE analysis as follows. Cells transformed with the relevant expression vector by electroporation were spread from a frozen glycerol stock onto an LB agar plate containing 0.2% (w/v) glucose and 25 μg/mL kanamycin and incubated at 30 °C overnight. The cells were then scraped into a small volume of LB medium and used to inoculate 3 mL of the relevant growth medium containing 25 μg/mL kanamycin in a 10 mL round bottom tube at a starting OD_600_ of 0.05. The culture was incubated at 37 °C with shaking at 220 rpm until the OD_600_ reached 0.5. One milliliter aliquots were transferred to clean round bottom tubes and isopropyl β-D-1-thiogalactopyranoside (IPTG) was added to a final concentration of 0.5 mM. The induced cultures were incubated with shaking at 220 rpm at 17 °C overnight or 30 °C for 3 h. A 500 μL aliquot of each culture was resuspended in lysis buffer (20 mM Tris, 0.5 M NaCl, 1% (v/v) Triton X-100, pH 8.0) and incubated at room temperature for 10 min. The cell lysate was centrifuged at 21,000 × *g* for 5 min (4 °C). The soluble fraction of the cell lysate was transferred to a tube containing 30 μL cOMPLETE His-Tag purification Ni-NTA resin (Roche, Basel, Switzerland) suspended in 500 μL buffer A (8 M urea, 20 mM Tris, 0.5 M NaCl, pH 8.0), while the insoluble fraction of the cell lysate was dissolved in 500 μL buffer A, centrifuged at 21,000 × *g* for 5 min, and then transferred to a tube containing 30 μL Ni-NTA resin suspended in 500 μL buffer A. In both cases, the resin was incubated at room temperature for 10 min, washed twice with 500 μL buffer A, and then eluted by incubation with 50 μL buffer B (8 M urea, 20 mM Tris, 0.5 M NaCl, 0.5 M imidazole, pH 8.0) at room temperature for 5 min. Fifteen microliters of supernatant was mixed with 5 μL of 4× SDS-PAGE sample loading buffer and heated at 90 °C for 10 min, then loaded onto a 4–15% pre-cast SDS-PAGE gel (Bio-Rad, Hercules, CA). The gel was run at 200 V for 30 min and visualized with Coomassie Blue.

### Large-scale protein expression and purification

For expression and purification of the ‘*Ca*. P. ubique’ SBPs, *E. coli* BL21(DE3) or SHuffle T7 cells transformed with the relevant expression vector were spread from a frozen glycerol stock onto an LB agar plate containing 0.2% (w/v) glucose and 25 μg/mL kanamycin, and incubated at 30 °C overnight. The cells were then scraped into 3 mL LB medium, and 500 μL of the resulting cell suspension was used to inoculate 500 mL LB or TB medium supplemented with 25 μg/mL kanamycin in a 2 L or 3 L flask, preheated at 37 °C. The culture was incubated at 37 °C with shaking at 220 rpm until the OD_600_ reached 0.5, then cooled briefly in an ice-water bath until the temperature reached ~25 °C. IPTG was added to a concentration of 0.5 mM, and the culture was incubated at 17 °C with shaking at 220 rpm for a further 16 h. Cells were pelleted by centrifugation (3300 × *g*, 15 min, 4 °C) and frozen at −20 °C until use. For protein purification, cells were thawed on ice, resuspended in 100 mL Ni binding buffer (20 mM Tris, 500 mM NaCl, 20 mM imidazole, pH 8.0), and lysed by sonication. After addition of 500 U Benzonase Nuclease (Sigma-Aldrich, St. Louis, MO) to digest DNA, the cell lysate was centrifuged at 10,000 × *g* for 1 h (4 °C). The supernatant was filtered through a 0.45 μm syringe filter and then loaded onto a 1 mL HisTrap HP column (Cytiva, Marlborough, MA) equilibrated with Ni wash buffer using an ÄKTA Pure FPLC system (Cytiva). For purification under native conditions, the column was washed with 10 mL Ni binding buffer followed by 10 mL Ni wash buffer (20 mM Tris, 500 mM NaCl, 44 mM imidazole, pH 8.0), and then the target protein was eluted in 10 mL Ni elution buffer (20 mM Tris, 500 mM NaCl, 500 mM imidazole, pH 8.0). For purification under denaturing conditions, the column was washed with denaturing Ni binding buffer (8 M urea, 20 mM Tris, 250 mM NaCl, 20 mM imidazole, pH 8.0) at 1 mL/min for 30 min after loading of the clarified cell lysate, and the target protein was eluted with 10 mL denaturing Ni elution buffer (8 M urea, 20 mM Tris, 250 mM NaCl, 250 mM imidazole, pH 8.0). Proteins purified under native conditions were concentrated to 400 μL using a 10 kDa molecular weight cutoff (MWCO) Amicon Ultra-4 centrifugal spin concentrator (Merck-Millipore, Burlington, MA) and purified by size-exclusion chromatography using a Superdex 200 Increase 10/300 column (Cytiva), eluting in DSF buffer (20 mM HEPES, 0.3 M NaCl, pH 7.50). For storage, proteins were concentrated to a volume of 0.5–2 mL and glycerol was added to a concentration of 10% (v/v). The protein was then flash-frozen in 100–200 μL aliquots in liquid nitrogen and stored at −80 C until use. ArgT from *Salmonella enterica* was expressed from a pETMCSIII plasmid and purified as described previously^58^.

### Protein refolding

Protein purified under denaturing conditions was diluted to a concentration of 0.5 mg/mL and volume of 10–30 mL in denaturing Ni binding buffer (8 M urea, 20 mM Tris, 250 mM NaCl, 20 mM imidazole, pH 8.0) and transferred to 10 kDa MWCO SnakeSkin dialysis tubing (Thermo Scientific, Waltham, MA). The protein was then dialyzed against 2 L dialysis buffer (20 mM Tris, 150 mM NaCl, pH 8.0) at 4 °C with three buffer changes over a period of 24 h. The protein was collected and exchanged into DSF buffer using a 10 kDa MWCO Amicon Ultra-15 centrifugal concentrator, then concentrated to 400 μL and purified by size-exclusion chromatography as described above.

### Differential scanning fluorimetry

DSF experiments were performed using a StepOnePlus Real-Time PCR System (Applied Biosystems, Bedford, MA) based on literature protocols^59,60^. Reaction mixtures were prepared in twin.tec Real-Time PCR Plates (Eppendorf, Hamburg, Germany) and contained 5× SYPRO Orange (Sigma-Aldrich), 2.5 μM protein, and 2 μL 10× ligand in a total volume of 20 μL DSF buffer. The plate was sealed with optically clear sealing film and centrifuged at 2000 × *g* for 1 min before loading into the real-time PCR instrument. The temperature was ramped at a rate of 1% (approximately 1.33 °C/min), typically over a 60 °C window centered on the melting temperature (*T*_M_) of the target protein. Fluorescence was monitored using the ROX channel. *T*_M_ values were determined by taking the derivative of fluorescence intensity with respect to temperature and fitting the resulting data to a quadratic equation in a 6 °C window in the vicinity of the *T*_M_ in R software.

Proteins were initially screened for binding to metabolites in four Phenotype MicroArray plates, PM1 to PM4, from Biolog (Hayward, CA). The contents of each well were dissolved in 50 μL (PM1 to PM3) or 20 μL (PM4) sterile filtered water, giving a concentration of approximately 10-20 mM in each well^60^. The plates were then sealed with aluminium sealing films and stored at −80 °C. Prior to use, the plates were thawed at room temperature and then shaken at 30 °C until the compounds had redissolved. Two microliters of each compound was added to 18 μL reaction mixture prepared as described above. A 2 °C increase in *T*_M_ compared with the median value across the plate was taken as indicative of binding^60^.

For screening of individual compounds and confirmatory assays, compounds were dissolved at a concentration of 100 mM in ligand buffer (0.1 M HEPES pH 7.5), and the pH was adjusted with 1 M NaOH or 1 M HCl if necessary (specifically, if the pH of a 10 mM solution of the compound diluted in DSF buffer fell outside the range 6.5–8.0). These stock solutions were stored at −20 °C. Two microliters of each compound was directly added to 18 μL reaction mixture, giving a final concentration of 10 mM, or first diluted 10-fold or 100-fold in DSF buffer to give final concentrations of 1 mM or 0.1 mM in the assay. A list of chemicals used for screening, including the supplier and catalog number, is provided in **Table S3**.

### Isothermal titration calorimetry

ITC experiments were performed using a MicroCal PEAQ-ITC system (Malvern Panalytical, Malvern, United Kingdom). Protein samples were refolded and freshly purified (not frozen), and protein and ligand samples were prepared in the same batch of DSF buffer used for size-exclusion chromatography to minimize the heat of dilution. Experiments were performed at 25 °C with stirring at 700 rpm and 10 μcal/s reference power. Titration parameters were varied depending on the protein yield, the fraction of active protein, and the affinity and enthalpy of the interaction. In a typical titration, 35 μM protein was titrated with 1 × 0.4 μL and 19 × 1.6 μL injections of ligand, with the ligand concentration chosen to give >1.5-fold molar excess of ligand to active protein at the end of the titration. ITC experiments were generally performed at least in duplicate.

For simple 1:1 binding interactions, the association constant (*K*_a_), enthalpy (Δ*H*), and stoichiometry (*n*) of the interaction were determined by fitting the data to the one-set-of-sites model in MicroCal PEAQ-ITC analysis software. In the case of the SAR11_0769 + D-glucose interaction, thermodynamic parameters were estimated through Bayesian fitting to a modified competitive binding model, which incorporated an additional parameter to account for the fraction of the ligand in each anomeric form, and a two-sets-of-sites model implemented in pytc software^61^; the latter model is equivalent to the two-sets-of-sites model in the MicroCal software, except without the minor correction for heat associated with the displaced volume for each injection (for consistency with the other models in pytc). Thermodynamic parameters for the SAR11_0953 + L-glutamate, SAR11_1210 + L arginine, and SAR11_1336 + glycine betaine interactions were determined through competitive displacement experiments^62^, in which L-phenylalanine, D-octopine, or glycine (respectively) were included at a fixed concentration in the cell to reduce the apparent binding affinity for the ligand of interest. The data for these competitive binding experiments was analyzed by Bayesian fitting to the competitive binding sites model in pytc software. Finally, to confirm the high affinity of the SAR11_1210 + L-arginine interaction, a competitive binding experiment was performed where SAR11_1210 and ArgT from *Salmonella enterica* (which has a *K*_d_ of 15 nM for L-arginine) were included in the cell together at the same concentration (28 μM) and titrated with L-arginine. The data was fitted to the two-sets-of-sites binding model as described above to obtain thermodynamic parameters for both interactions. For all analyses, the heat of dilution was assumed to be a small constant value and included as a fitted parameter in the model. The validity of this assumption was confirmed for each ligand by performing a control titration where the ligand was injected into DSF buffer.

### Spectrophotometric analysis of iron(III) binding

Binding of iron(III) to SAR11_1238 was analyzed using a spectrophotometric assay based on literature protocols^63,64^. UV-vis spectra were recorded at room temperature (25 °C) in a 96-well plate from 300 nm to 630 nm with 1 nm bandwidth using a Multiskan GO spectrophotometer (Thermo Scientific). An initial protein concentration of 100 μM and an initial volume of 200 μL were used for all spectrophotometric assays. Firstly, purified SAR11_1238 was thawed and exchanged into 50 mM Tris, 200 mM NaCl buffer (pH 8.0) using a centrifugal concentrator, and the spectrum of the resulting protein sample was recorded. To prepare unliganded protein for iron-binding assays, the protein was exchanged into 50 mM Tris, 200 mM NaCl, 20 mM sodium citrate buffer (pH 8.0) by three rounds of 30-fold dilution and concentration, allowing chelation and removal of the metal ligand. Citrate was then removed by four rounds of 30-fold dilution and concentration with 50 mM Tris, 200 mM NaCl buffer (pH 8.0). Binding assays were performed by titrating the unliganded protein (200 μL of 100 μM solution) with eight or ten 5 μL injections of 800 μM iron(III) solution, which was prepared from iron(III) chloride and a 2.5-fold molar excess of trisodium citrate (which ensures that the iron(III) remains soluble) in ultrapure water. To confirm that SAR11_1238 binds iron(III) rather than the iron(III)-citrate complex, the protein was also titrated under the same conditions with 800 μM ammonium iron(II) sulfate; under the aerobic conditions of the assay, iron(II) is rapidly oxidized to iron(III) ^63^. UV-vis spectra were recorded 1 min (iron(II)) or 15 min (iron(III)) after each injection. Finally, a competitive binding assay with citrate was used to estimate the affinity of SAR11_1238 for iron(III). The protein was saturated with a 2-fold molar excess of iron(III) solution, diluted to a volume of 1 mL, and then dialyzed against 500 mL of 50 mM Tris, 200 mM NaCl buffer (pH 8.0) at 4 °C overnight to remove excess iron(III) and citrate. The protein was then concentrated to 100 μM and titrated with 5 μL injections of eight 2-fold serial dilutions of 500 mM sodium citrate (adjusted to pH 8.0 in 50 mM Tris, 200 mM NaCl buffer). The absorbance at 440 nm was recorded 5 min after each addition. The data were fitted to a hyperbolic curve, yielding an apparent *K*_d_ of 9.0 mM for citrate. Given that citrate has a *K*_d_ of ~10^−17^ M for iron(III), this implies that SAR11_1238 has a *K*_d_ for iron(III) on the order of ~10^−19^ M, similar to previously characterized iron(III)-binding proteins^64,65^.

### X-ray crystallography

SAR11_0769 and SAR11_1210 were expressed and purified by nickel affinity chromatography under native conditions as described above. After addition of a 20-fold molar excess of D-glucose (SAR11_0769) or L-arginine (SAR11_1210), the protein was purified further by size-exclusion chromatography on a HiLoad 26/600 Superdex 75 pg column (Cytiva), eluting in 3× crystallization buffer (60 mM HEPES, 150 mM NaCl, pH 7.5). Fractions containing the target protein were collected, and D-glucose (SAR11_0769) or L-arginine (SAR11_1210) was added to a concentration of 30 μM. The protein was concentrated to a volume of ~500 μL, diluted three-fold in water to reduce the NaCl concentration to 50 mM, and then concentrated further to 12 mg/mL. Protein crystallization was performed using the vapor diffusion method at 20 °C. The crystal used for structure determination of SAR11_0769 was obtained from a hanging drop containing 1 μL of 12 mg/mL protein and 1 μL of 18.5% (w/v) polyethylene glycol (PEG) 1500, 10% (v/v) isopropanol, 0.1 M Bis-Tris (pH 6.5) as the precipitant. The crystal was cryoprotected in 15% (w/v) PEG 1500, 30% (v/v) ethylene glycol, 10% (v/v) isopropanol, 0.1 M Bis-Tris (pH 6.5) and flash frozen in liquid nitrogen. The crystal used for structure determination of SAR11_1210 was obtained by serial microseeding from a crystal grown in a hanging drop containing 1 μL of 12 mg/mL protein and 1 μL of 21% (w/v) PEG 1500, 0.1 M MES (pH 6.0) as the precipitant. The crystal was cryoprotected in 30% (w/v) PEG 1500, 0.1 M MES (pH 6.0) and flash frozen in liquid nitrogen. X-ray diffraction data were collected on beamline BL32XU at the SPring-8 synchrotron (Harima, Japan), using the ZOO suite for automated data collection^66^. The data were automatically indexed, integrated, scaled and merged in XDS^67^ using KAMO^68^. The structure was solved by molecular replacement in Phaser (SAR11_0769, using a model generated in ColabFold as a search model) or MOLREP (SAR11_1210, using an opine-binding protein from *Agrobacterium fabrum*, PDB ID 5OT8, as a search model)^69–71^. The structures were then refined by iterative real-space and reciprocal-space refinement in REFMAC^72^, Phenix^73^, and COOT^74^. Simulated annealing in Phenix was performed as the first step in refinement of the SAR11_0769 structure to reduce model bias. Data collection and refinement statistics are given in **Table S9**.

### Biogeographical analysis

Abundance data for each SBP gene from ‘*Ca*. P. ubique’ HTCC1062 in the *Tara* Oceans OM-RGC_v2_metaG and OM-RGC_v2_metaT datasets was obtained through a BLAST search of the Ocean Gene Atlas v2.0 server^33^ with a stringent e-value threshold of 10^−40^. Plots were generated using GraphPad Prism and R.

### Phylogenetic analysis

Protein sequences homologous to the SBP of interest were identified *via* a BLAST search of the UniProtKB Reference Proteomes and Swiss-Prot databases^75^. The resulting sequences were filtered to remove a small number of unusually long sequences (>20% greater than mean length) and aligned in MUSCLE v3.8.31^76^. The alignment was trimmed in trimAl v1.2 using the automated1 option^77^ and then used to generate a maximum-likelihood phylogeny in FastTree v2.1.11, using LG + Γ_20_ as the substitution model^78^. For each protein sequence in the tree, the fraction of conserved binding site residues, compared with the corresponding protein from ‘*Ca*. P. ubique’ HTCC1062, was estimated. The binding site residues were obtained from the crystal structure (SAR11_0769) or estimated from an AlphaFold2 model^79,80^. For this analysis, the following substitutions were treated as conservative: S/T, I/M, V/L, I/V, L/M, D/E, Q/N, A/V, F/Y, Y/W, F/W. Figures were generated using the ggtree package in R^81^.

## Acknowledgments

B.E.C. was supported by a JSPS Postdoctoral Fellowship for Overseas Researchers from the Japan Society for the Promotion of Science. P.L. gratefully acknowledges funding from the Okinawa Institute of Science and Technology. We thank Assaf Vardi for critical reading of the manuscript, Dan Kozome for technical assistance with X-ray data collection, and the Instrumental Analysis Section at OIST for providing instrument access.

## Author contributions

Conceptualization: BEC, PL; Investigation: BEC; Analysis, BEC, UA, CJJ, PL; Supervision, PL; Writing – original draft: BEC; Writing – review and editing: BEC, UA, CJJ, PL.

## Data and materials availability

Coordinates and structure factors for the crystal structures of SAR11_0769 and SAR11_1210 have been deposited in the Protein Data Bank under accession codes 8HQQ and 8HQR, respectively. Raw ITC data, raw DSF data for Biolog assays, phylogenetic and biogeographical data, code, and source data for figures will be made available *via* the Open Science Framework prior to publication.

