## Supplementary Information for "Ultrahigh-affinity transport proteins from ubiquitous marine bacteria reveal mechanisms and global patterns of nutrient uptake"

**Supplementary Methods**

Refolding of SAR11_0271 and SAR11_1346

Firstly, SAR11_0271 and SAR11_1346 were purified under denaturing conditions from inclusion bodies. Cell pellets from 1 L cultures were thawed and resuspended in 100 mL IB lysis buffer (50 mM Tris, 300 mM NaCl, 10 mM DTT, pH 8.0). The cells were lysed by addition of Triton X-100, dropwise with stirring, to a final concentration of 0.5% (v/v), followed by incubation on ice for 15 min. The lysate was centrifuged at 10000 × g for 20 min, then the pellet was washed with 25 mL IB lysis buffer + 1% Triton X-100, 25 mL IB wash buffer (50 mM Tris, 1 M NaCl, 10 mM DTT, pH 8.0), and 25 mL IB lysis buffer. The pellet was dissolved by incubation overnight in 6 M guanidine HCl, 20 mM Tris, 300 mM NaCl, 1 mM DTT pH 8.0. The resulting suspension was centrifuged at 10000 × g for 20 min, filtered through a 0.45 µm syringe filter, and loaded onto a 1 mL HisTrap column equilibrated with denaturing Ni wash buffer. The column was washed with 15 mL denaturing Ni wash buffer, and the target protein was eluted with 5 mL denaturing elution buffer, then concentrated to 5 mg/mL using a 10 K MWCO centrifugal concentrator.

Next, small-scale refolding trials were performed to identify promising conditions for refolding. Twelve buffers were tested: 150 mM NaCl, 20 mM Tris pH 8.0 or 20 mM HEPES pH 7.0, with or without redox cycling (1 mM reduced glutathione + 0.1 mM oxidized glutathione), with or without an additive (5% glycerol or 0.4 M l-arginine). 4 µL of 5 mg/mL solubilized protein was diluted into 72 µL refolding buffer and incubated at 4 °C overnight. The soluble fractions of the resulting mixtures were loaded onto an SDS-PAGE gel (alongside the denatured starting material as a loading control) to identify conditions that gave a high yield of soluble protein. Both SAR11_0271 and SAR11_1346 showed a high level of soluble protein across all conditions.

Finally, the optimal refolding conditions identified in the small-scale refolding trials were scaled up to a 1.5–3.5 mg scale. SAR11_0271 and SAR11_1346 were diluted 1/25 from a 5 mg/mL stock into 20 mM Tris pH 8.0, 150 mM NaCl, 0.4 M L-arginine (SAR11_0271) or 20 mM Tris pH 8.0, 150 mM NaCl (SAR11_1346), and incubated at 4 °C overnight with gentle stirring. The proteins were filtered through a 0.45 µm syringe filter, concentrated to 400 µL, loaded onto a Superdex 200 Increase 10/300 column, and eluted in DSF buffer. However, only high-molecular-weight protein aggregates were observed for both proteins.

Systematic search for affinity

To identify $K_d$ values of previously characterized bacterial and archaeal SBPs, searches of Pubmed and Web of Knowledge were conducted with the search terms given below. Synthetic constructs such as biosensors were excluded. Although these searches do not necessarily encompass every SBP reported in the literature, the resulting SBPs constitute a large and representative subset of those reported.

*Sugars*:
1. ("sugar-binding protein" OR "glucose-binding protein" OR "oligosaccharide-binding protein" OR "monosaccharide-binding protein" OR "disaccharide-binding protein") **AND** ("affinity" OR "Kd" OR "dissociation constant")
2. ("glucose" OR "sugar" OR "oligosaccharide" OR "monosaccharide" OR "disaccharide") AND ("solute-binding protein" OR "substrate-binding protein" OR "periplasmic-binding protein" OR "ABC transporter" OR "ABC transport protein" OR "ATP-binding cassette") **AND** ("affinity" OR "Kd" OR "dissociation constant")

*Amino acids:*
1. ("amino acid" OR "alanine" OR "cysteine" OR "aspartate" OR "glutamate" OR "phenylalanine" OR "glycine" OR "histidine" OR "isoleucine" OR "lysine" OR "leucine" OR "methionine" OR "asparagine" OR "proline" OR "glutamine" OR "arginine" OR "serine" OR "threonine" OR "tryptophan" OR "tyrosine" OR "pyroglutamate" OR "cystine" OR "glutathione" OR "glutamic acid" OR "aspartic acid") **AND** ("solute-binding protein" OR "substrate-binding protein" OR "periplasmic-binding protein" OR "ABC transporter" OR "ABC transport protein" OR "ATP-binding cassette") **AND** ("affinity" OR "Kd" OR "dissociation constant")
2. ("amino acid-binding protein" OR "alanine-binding protein" OR "aspartate-binding protein" OR "glutamate-binding protein" OR "glycine-binding protein" OR "histidine-binding protein" OR "lysine-binding protein" OR "leucine-binding protein" OR "methionine-binding protein" OR "proline-binding protein" OR "glutamine-binding protein" OR "arginine-binding protein" OR "threonine-binding protein" OR "tryptophan-binding protein" OR "tyrosine-binding protein" OR "cystine-binding protein" OR "glutathione-binding protein" OR "glutamic acid-binding protein" OR "amino acid-binding proteins") **AND** ("affinity" OR "Kd" OR "dissociation constant")

*Osmolytes and related compounds:*
1. ("quaternary amine" OR "osmolyte" OR "compatible solute" OR "osmoprotectant" OR "quaternary amines" OR "osmolytes" OR "compatible solutes" OR "osmoprotectants" OR "betaine" OR "choline" OR "ectoine" OR "DMSP" or "glycine" OR "carnitine" OR "trimethylamine-N-oxide" OR "taurine" OR "dimethylsulfoniopropionate" OR "TMAO") **AND** ("solute-binding protein" OR "substrate-binding protein" OR "periplasmic-binding protein" OR "ABC transporter" OR "ABC transport protein" OR "ATP-binding cassette") **AND** ("affinity" OR "Kd" OR "dissociation constant")
2. ("ChoX" OR "ProX" OR "OpuAC" or "OpuBC" or "OpuC") **AND** ("solute-binding protein" OR "substrate-binding protein" OR "periplasmic-binding protein" OR "ABC transporter" OR "ABC transport protein" OR "ATP-binding cassette") **AND** ("affinity" OR "Kd" OR "dissociation constant")

*General*

(("solute-binding protein" OR "substrate-binding protein" OR "periplasmic-binding protein" OR "ABC transporter" OR "ABC transport protein" OR "ATP-binding cassette") **AND** ("affinity" OR "Kd" OR "dissociation constant" OR "association constant" OR "Ka")) **NOT** (P-glycoprotein)

### Supplementary Text

*Supplementary Note 1. Explanation of discrepancies between DSF and ITC data.*

In the case of SAR11_0797 (annotated as a glycine betaine-binding protein), no significant heat was observed when the protein was titrated with choline-*O*-sulfate or phosphocholine, which gave the highest $\Delta T_M$ values in DSF experiments (8.0 °C and 6.6 °C, respectively). Because we performed *in vitro* protein refolding prior to ITC, due to the known propensity of SBPs to co-purify with endogenously bound ligands[1], we considered the possibility that the refolded protein was misfolded and inactive, and repeated the titrations with natively purified protein; however, binding was still not observed. The $\Delta T_M$ values observed for SAR11_0797 with choline-*O*-sulfate and phosphocholine are relatively low compared with the other SBPs (**Fig. 2**), suggesting that these metabolites might be structurally similar but not identical to the physiological ligand(s). Given the known capability of the SAR11 bacterium '*Candidatus* Pelagibacter giovannonii' to transport and metabolize sulfonates[2], which are structurally similar to choline-*O*-sulfate, we considered the possibility that SAR11_0797 is a sulfonate-binding protein, but binding of physiologically important compounds such as taurine, 2,3-dihydroxypropane-1-sulfonate, and isethionate (2-hydroxyethane sulfonate) was not observed by DSF. Binding of choline, glycine betaine, proline, and proline betaine was also not observed.

Additionally, binding of SAR11_0655 to L-glutamine ($\Delta T_M$ 12.3 °C) and SAR11_0769 to cellobiose ($\Delta T_M$ 16.7 °C) and maltotriose ($\Delta T_M$ 20.1 °C) could only be detected by DSF. The interaction between SAR11_0655 and L-glutamine may have been obscured by a binding enthalpy close to zero, given the low positive enthalpy of binding observed for the confirmed ligand L-pyroglutamate ($\Delta H$ = +4.5 kcal/mol). In the case of SAR11_0769, contamination of the cellobiose and maltotriose solutions with trace amounts of D-glucose (a high-affinity ligand of SAR11_0769 and the monomeric unit of the oligosaccharides cellobiose and maltotriose) is likely responsible for the false positives in DSF.

*Supplementary Note 2. Binding mode of L-arginine to SAR11_1210.*

SAR11_1210 belongs to the polar amino acid-binding protein (AABP) family of SBPs, which are composed of two α/β domains connected by two hinge strands (**Fig. S6A**). Ligand binding in SBPs is typically accompanied by a conformational change from an open conformation, with a large open cavity between the two α/β domains, to a closed conformation, with the two α/β domains encapsulating the ligand at their interface. Thus, binding affinity in SBPs is governed not only by the strength of molecular interactions between the protein and ligand, but also the intrinsic conformational equilibrium between the open and closed conformational states of the protein[3]. Indeed, mutations in the hinge region of SBPs are known to affect binding affinity *via* their effect on this conformational equilibrium[3–5]. The binding pockets of AABPs typically have a conserved region containing the binding motif for the amino acid moiety and a variable region that accommodates various amino acid side chains. The canonical binding mode for L-arginine binding in SBPs is exemplified by the structure of lysine/arginine/ornithine-binding protein from

*Geobacillus stearothermophilus* (PDB 2Q2A): the amino acid moiety is bound by the side chains of Thr90, Arg95, and Asp200, and the backbone amide groups of Gly88, Thr90 and Thr161, while the side chain forms π-stacking interactions with Phe32 and Trp70, and polar interactions with Asp29, Glu36, Ser87, and Gln157 (**Fig. S6D**). This binding mode is shared by most other L-arginine-binding SBPs in the Protein Data Bank (1LAF, 2Y7I, 3N26, 3VVF, 4H5F, 4I62, 4PSH, 4YMX, 4ZV1, and 5T0W, although not 3TQL or 6DET), with some minor variations of the residues that bind the guandinium group of the ligand.

In SAR11_1210, the binding motif for the guandinium group and the overall pose of L-arginine in the binding site are typical of L-arginine-binding SBPs (**Fig. S6B-C**). However, this protein shows an unusual variation of the typical amino acid binding motif in which a conserved interaction between Asp200 and the amino group of the ligand is replaced by an interaction with Glu108 in the hinge region, which usually does not contribute directly to ligand binding. The direct involvement of the flexible hinge region in ligand binding would likely contribute to stabilization of the closed conformation of the protein in the presence of ligand, leading to an indirect increase in binding affinity, providing a possible explanation for the increased affinity of SAR11_1210 compared with homologous L-arginine-binding SBPs.

*Supplementary Note 3. Binding mode of D-glucose to SAR11_0769.*

Titration of SAR11_0769 with D-glucose consistently yielded an unusual biphasic binding isotherm across three independent protein preparations. One potential explanation for the biphasic binding curve is that SAR11_0769 has two binding sites for glucose, which is plausible given that some SBPs have multiple monosaccharide binding sites to enable binding of oligosaccharides. Fitting the ITC data to a two-sets-of-sites binding model gave a total binding stoichiometry of 1.43 ± 0.10 (mean ± s.d., $n$ = 4), consistent with ~70% of the refolded protein showing binding activity, and an upper limit of $K_d$ of ~59 pM for the high-affinity site and ~74 nM for the low-affinity site (**Fig. S13A**). An alternative explanation for the biphasic binding curve is that SAR11_0769 shows differential binding to the α and β anomeric forms of glucose; in this scenario, the first phase would represent binding of both anomers until the protein reaches saturation, and the second phase would represent displacement of the low-affinity anomer from the binding site by the high-affinity anomer. Fitting the ITC data to a modified competitive binding model with an additional parameter to account for the anomeric ratio of the ligand gave an upper limit of $K_d$ of ~27 pM for the high-affinity anomer and ~36 nM for the low-affinity anomer (**Fig. S13B**). The apparent anomeric ratio was 0.46 ± 0.04, which presumably deviates slightly from the known ratio of 0.35–0.40 to 0.60–0.65 α:β due to slow interconversion of the anomers by mutarotation on the timescale of the experiment (~1 h). Both the two-sets-of-sites and competitive binding models gave an equally good fit to the ITC data; thus, to distinguish between these two possible binding modes, we solved a high-resolution (1.86 Å) crystal structure of SAR11_0769 complexed with glucose (**Fig. S13C-E**). Electron density for only one molecule of β-D-glucose in each subunit could be identified (**Fig. S13D**), supporting the presence of a single binding site and a difference in binding affinity between the α and β anomers.

*Supplementary Note 4. Presence of a TRAP transporter for L-pyroglutamate in SAR11 bacteria.*
An unpublished crystal structure of a TRAP SBP from the SAR116 Alphaproteobacterium HIMB100 bound to co-purified L-pyroglutamate (1.40 Å) has been deposited in the Protein Data Bank (PDB ID 6WGM, UniProt ID G5ZWD6; Fedorov *et al.*, 2020). TRAP SBPs that show high sequence identity and binding site conservation compared with this protein are found in several strains of '*Ca*. P. ubique' such as HTCC9565 (NCBI accession WP_169035489.1, 44.7% i.d.), although not HTCC1062. In general, the gene for this pyroglutamate TRAP SBP is more abundantly distributed and expressed than SAR11_0655 in the surface ocean, except at high latitudes (**Fig. S14**). Together, these results indicate that SBP-dependent transporters for L-pyroglutamate are found widely in SAR11 bacteria.

*Supplementary Note 5. Carbon sources of SAR11 bacteria.*
The range of potential carbon sources that can be utilized by SAR11 bacteria is not entirely clear[6]. Early analysis based on the genome sequence of '*Ca.* P. ubique' HTCC1062 suggested a capacity of SAR11 bacteria to use D-glucose as a carbon source, due to the presence of a variant of the Entner-Doudoroff pathway for glycolysis[7]. However, subsequent analysis showed that the capacity for glucose assimilation is not universal among SAR11 ecotypes[8], and pyruvate was instead hypothesized to represent the universal carbon source of SAR11 bacteria[6]. It is known that '*Ca*. P. ubique' HTCC1062 can use oxaloacetate, but not succinate or malate, as a source of pyruvate in defined medium[9], although these compounds are likely useful when supplied at lower, physiologically relevant concentrations in a balanced nutrient mixture found in natural environments. Our results support the hypothesis that gluconeogenic precursors of pyruvate are environmentally important, universal carbon sources for SAR11 bacteria and suggest that a range of dicarboxylates are imported to fulfil this requirement.

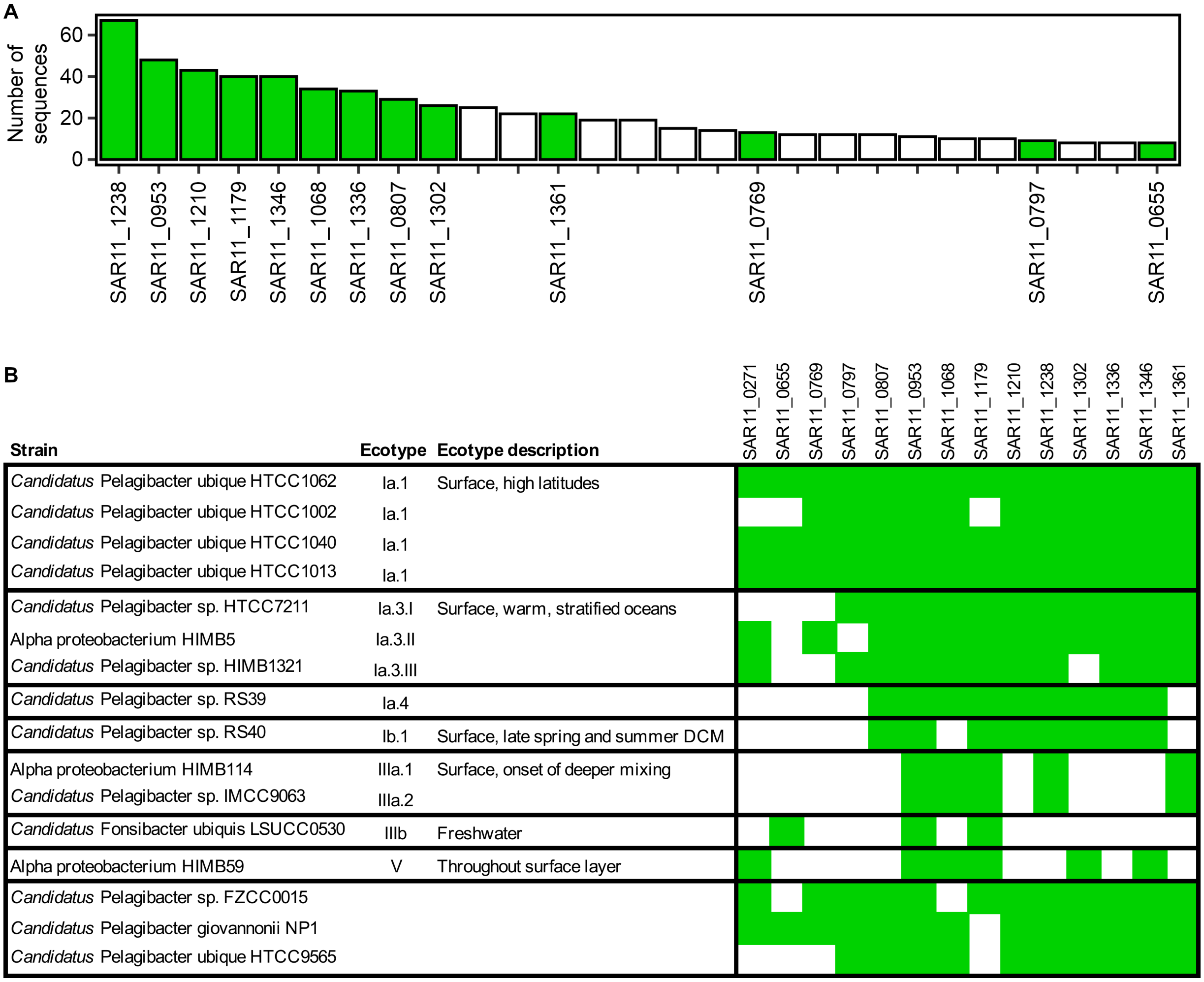


**Fig. S1. Distribution of '*Ca.* P. ubique' HTCC1062 SBPs among SAR11 bacteria.** (**A**) SBP sequences belonging to the SBP PFAM clans CL0177 and CL0144 from candidate order '*Pelagibacterales*' were obtained from the InterPro database and clustered at 35% sequence identity using mmseqs2 [10]. The graph shows the number of sequences in each ABC transporter-associated SBP cluster containing ≥8 sequences. Clusters that are represented in '*Ca.* P. ubique' HTCC1062 are shown in green. (**B**) Distribution of homologs of SBPs from '*Ca.* P. ubique' HTCC1062 in SAR11 strains with complete genome sequences. Homologs sharing <50% sequence identity with the corresponding protein in '*Ca*. P. ubique' HTCC1062 are not shown. Ecotype classifications are taken from ref. 11 and descriptions are taken verbatim from ref. 12.

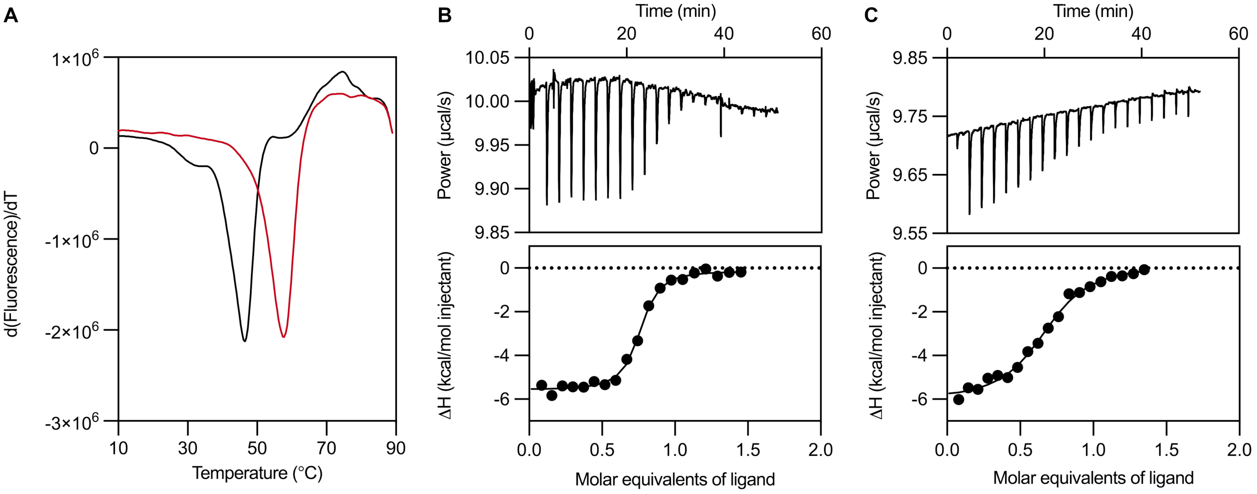


**Fig. S2. SAR11_1179 is a phosphate-binding protein.** (**A**) Thermal denaturation of SAR11_1179 in the absence (black) or presence (red) of 1 mM $Na_2HPO_4$ measured by DSF ($\Delta T_M$ = 10.9 ± 0.4 °C, mean ± s.d., $n$ = 3 technical replicates). (**B-C**) Representative ITC data for titration of 23 µM SAR11_1179 with 185 µM $Na_2HPO_4$ in the absence (**B**) or presence (**C**) of 28 mM $Na_2SO_4$. Fitting the data to the one-set-of-sites model gave a $K_d$ of 133 ± 28 nM in the absence of sulfate and 892 ± 122 nM in the presence of 28 mM sulfate (mean ± s.d., $n$ = 3 or 4 replicate titrations).

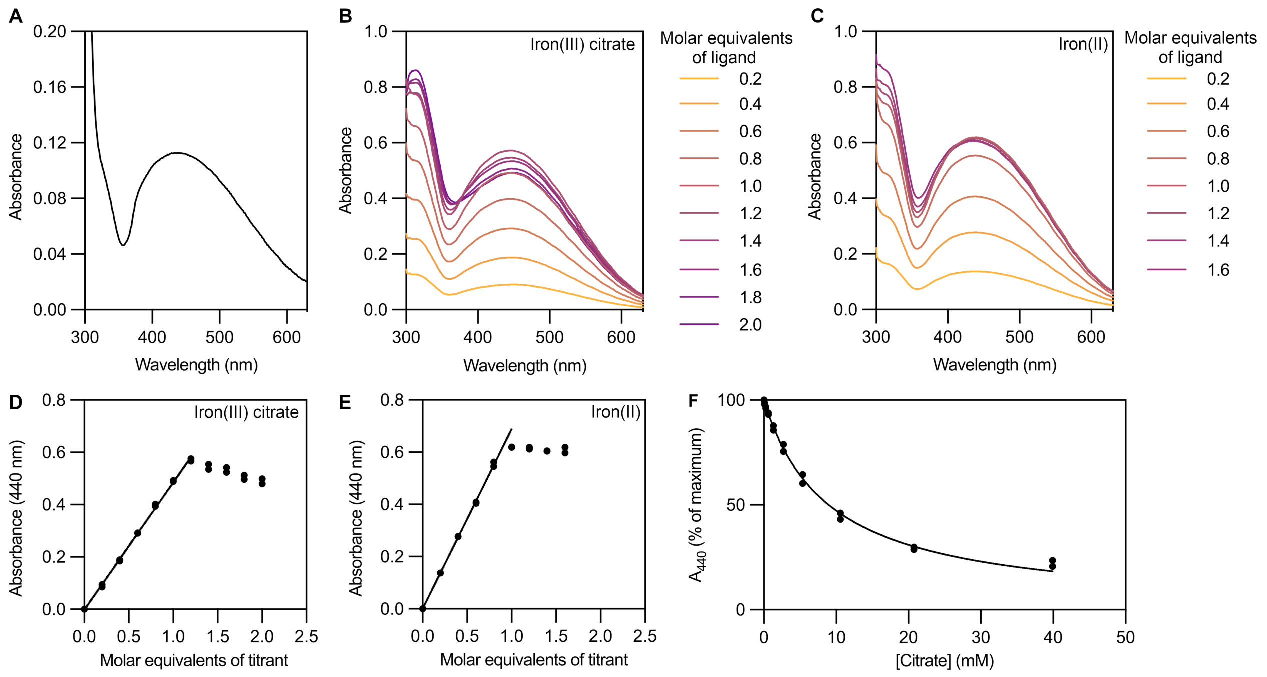


**Fig. S3. SAR11_1238 is an iron(III)-binding protein.** (**A**) UV-visible spectrum of SAR11_1238 purified from *E. coli*, without addition of ligand, showing presence of endogenously bound iron(III). (**B-C**) UV-visible spectra of unliganded SAR11_1238 titrated with iron(III) delivered as (**B**) iron(III) citrate or (**C**) ammonium iron(II) sulfate (see **Materials and Methods** for further explanation). (**D-E**), Titration of SAR11_1238 with iron(III) delivered as (**D**) iron(III) citrate or (**E**) ammonium iron(II) sulfate, monitored by absorbance at 440 nm. Discrete data points from four (D) or two (E) technical replicates (independent titrations) are shown. The line represents a fit to the linear portion of the titration. (**F**) Competitive titration of iron(III)-bound SAR11_1238 with citrate, monitored by absorbance at 440 nm. Results from two technical replicates (independent titrations) are shown as discrete data points.

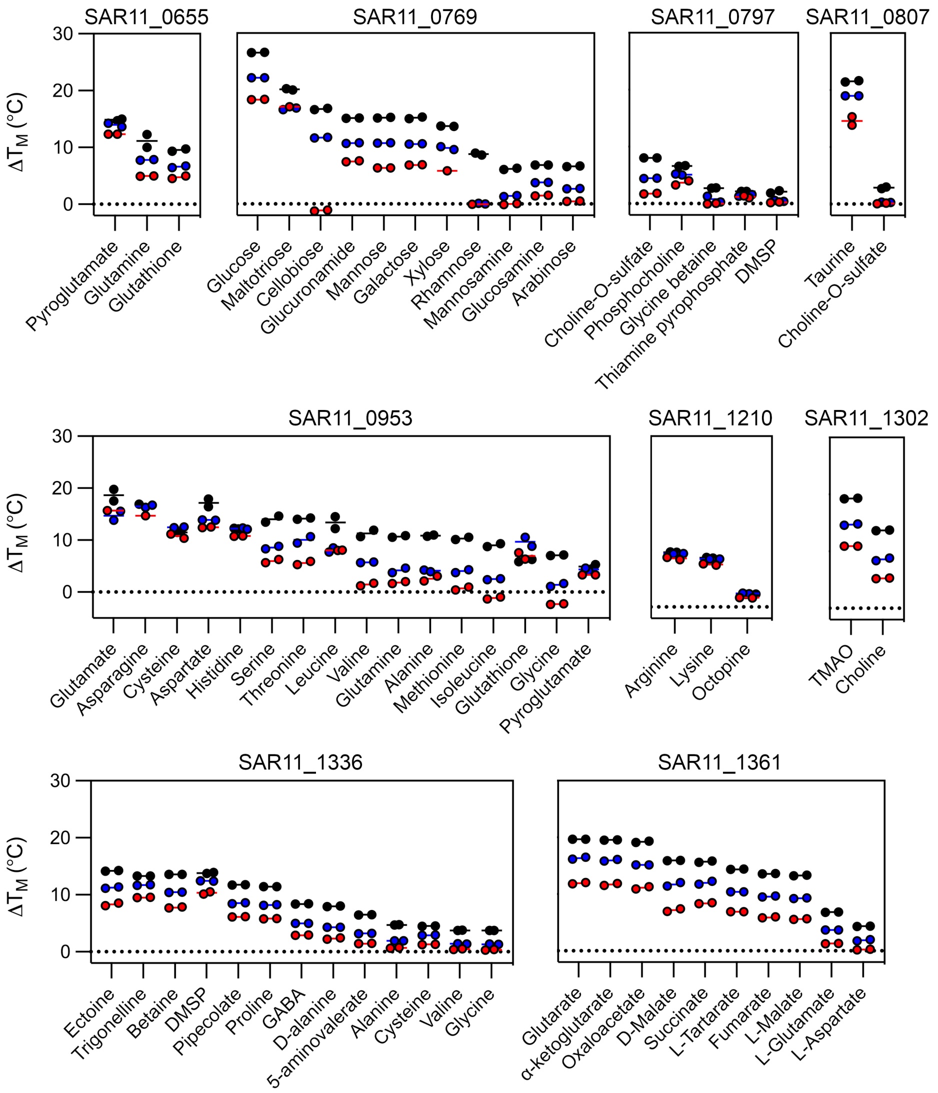


**Fig. S4. Concentration dependence of Δ$T_M$ values obtained by DSF.** For each SBP-ligand interaction, Δ$T_M$ was measured at three ligand concentrations: 10 mM (black), 1 mM (blue), and 0.1 mM (red). In DSF, specific ligands tend to show a steady decrease in Δ$T_M$ with decreasing concentration, as observed in most cases, whereas non-specific ligands tend to show a sudden drop in Δ$T_M$ with decreasing concentration[13]. Two technical replicates are shown for each condition. Data for 10 mM ligand is duplicated from **Fig. 2**. Abbreviations: DMSP, dimethylsulfoniopropionate; TMAO, trimethylamine-*N*-oxide; GABA, γ-aminobutyrate.

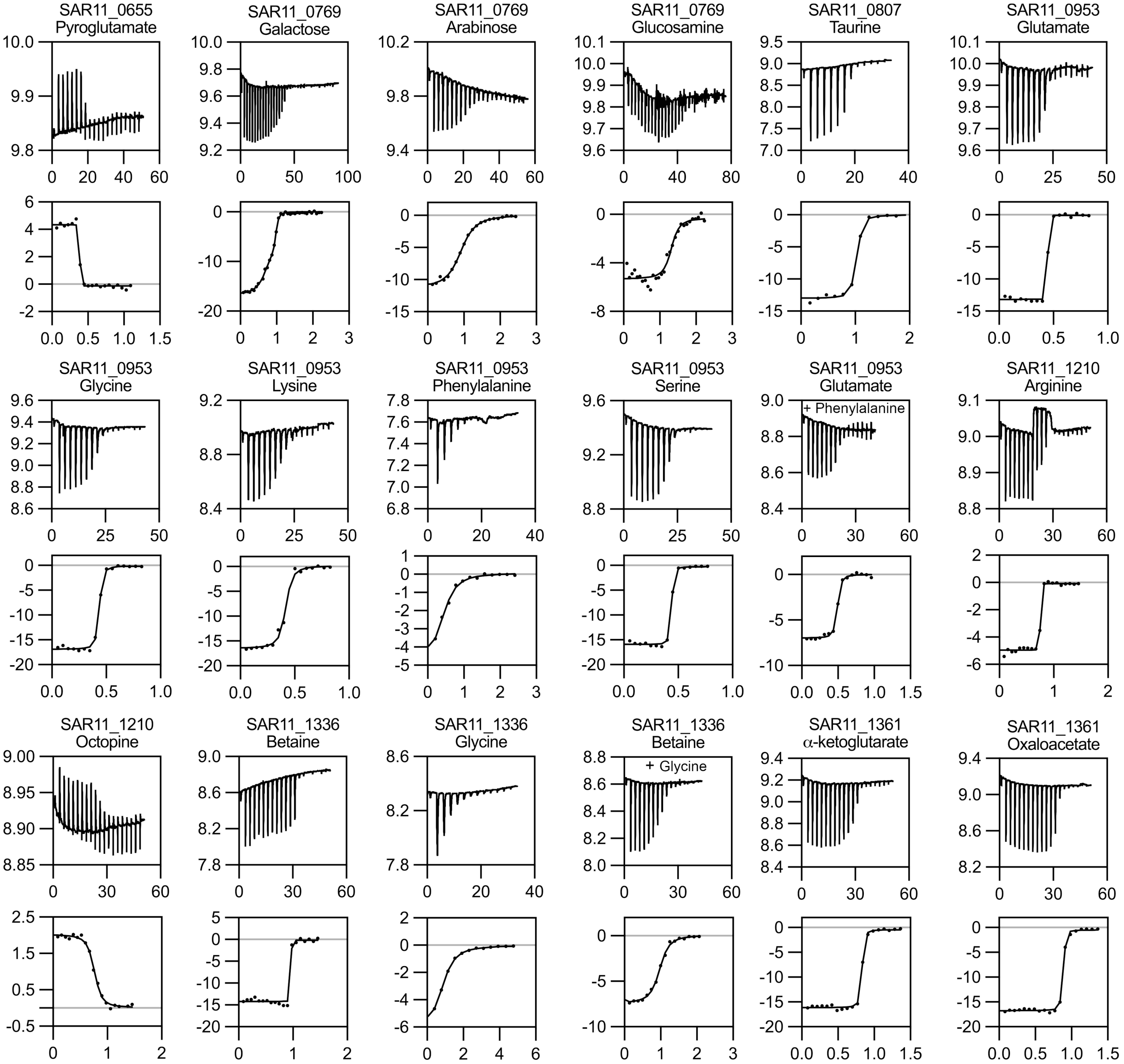


**Fig. S5. Representative ITC data for each SBP-ligand interaction.** For each SBP-ligand interaction, the top panel shows power (µcal/s) versus time (min) and the bottom panel shows $\Delta H$ (kcal/mol injectant) versus molar equivalents of ligand. Binding parameters are given in **table S5**. The remaining ITC data is shown in **Fig. 3** and **Fig. S2**.

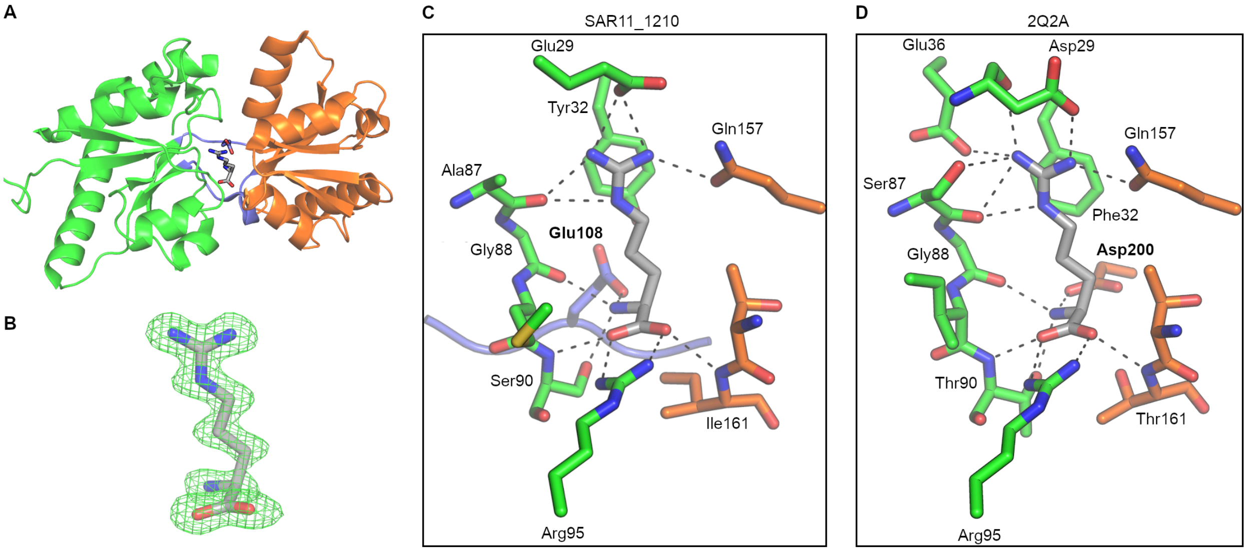


**Fig. S6. Binding mode of L-arginine to SAR11_1210.** (**A-C**) Crystal structure of SAR11_1210 complexed with L-arginine (1.32 Å). The large domain, small domain, and hinge regions are shown in green, orange, and purple, respectively. (**A**) Overall structure. (**B**) Electron density for the L-arginine molecule, shown by an $mF_o - dF_c$ map contoured at +3σ. (**C-D**) Comparison of binding modes of L-arginine to (**C**) SAR11_1210 and (**D**) a homologous lysine-/arginine-/ornithine-binding protein from *Geobacillus stearothermophilus* ($K_d$ 39 nM, PDB ID 2Q2A), which shows the amino acid binding motif typical of this SBP family[14]. Residues are numbered according to the homologous positions in SAR11_1210.

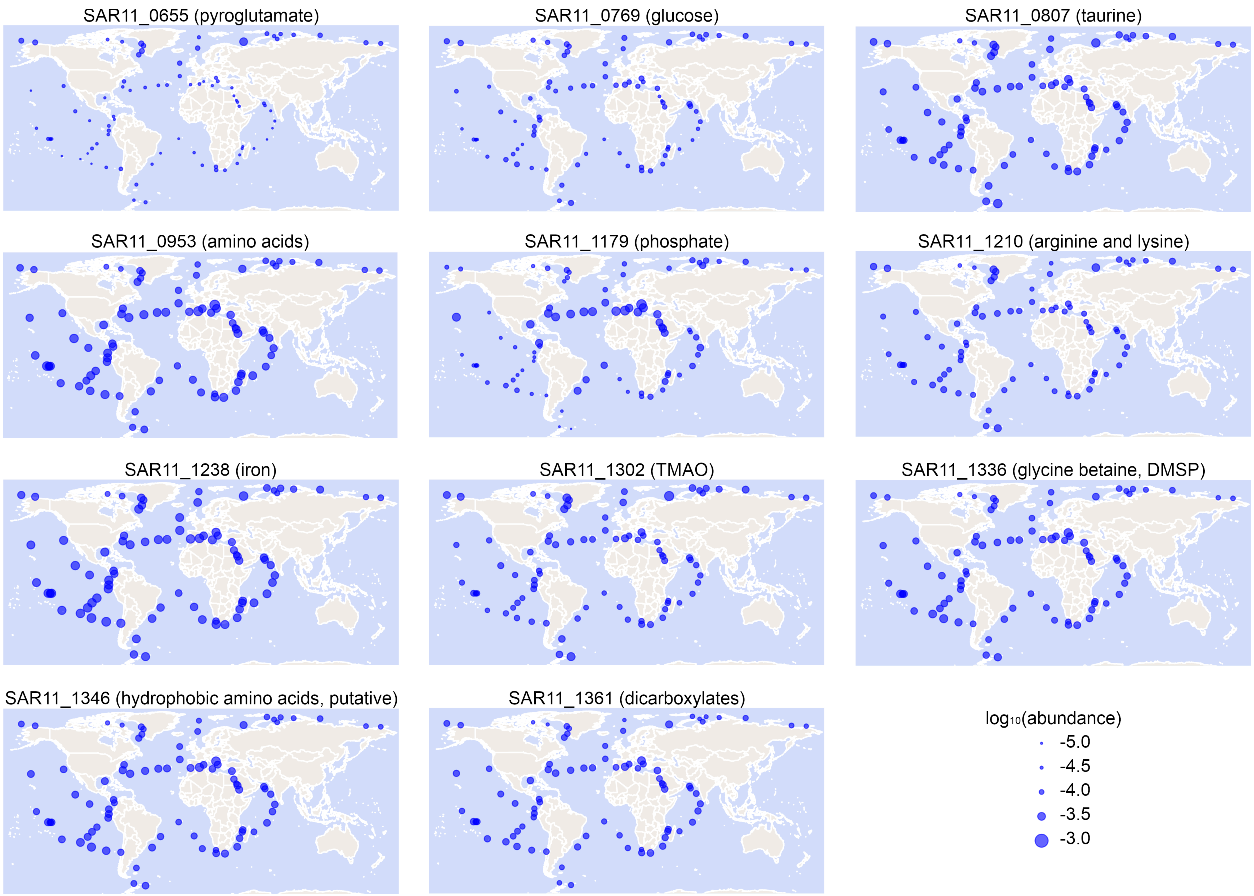


**Fig. S7. Abundance of SBP genes from '*Ca*. P. ubique' HTCC1062 in the surface ocean metagenome.** The abundance of each gene in surface samples from the *Tara* Oceans OM-RGCv2+G metagenomic dataset is shown. Data were obtained from the Ocean Gene Atlas v2.0 using an e-value cut-off of $10^{-40}$. Abundance at each location is expressed as the percentage of mapped reads and represented by point area on a linear scale.

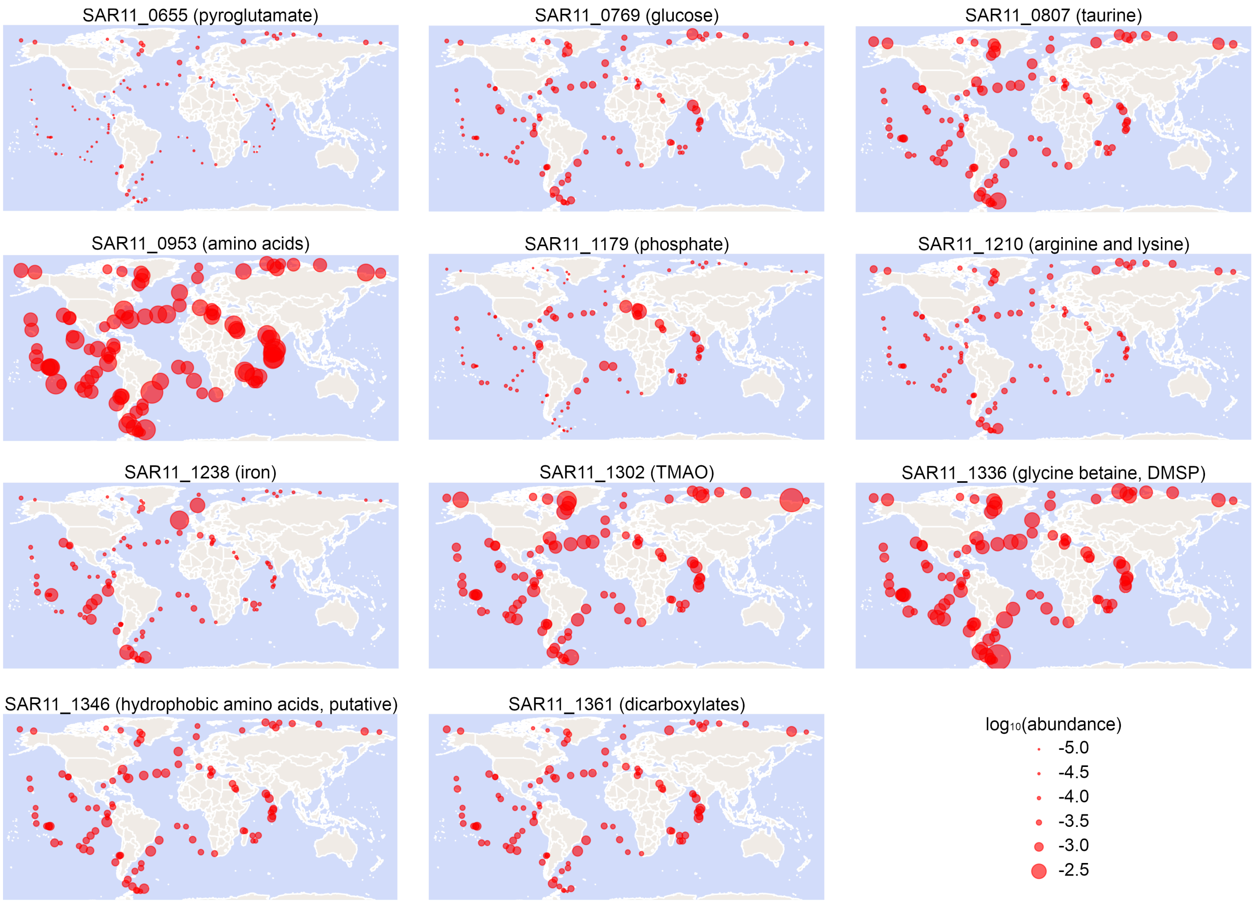


**Fig. S8. Abundance of SBP genes from '*Ca*. P. ubique' HTCC1062 in the surface ocean metatranscriptome.** The abundance of each gene in surface samples from the *Tara* Oceans OM-RGCv2+T metatranscriptomic dataset is shown. Data were obtained from the Ocean Gene Atlas v2.0 using an e-value cut-off of $10^{-40}$. Abundance at each location is expressed as the percentage of mapped reads and represented by point area on a linear scale.

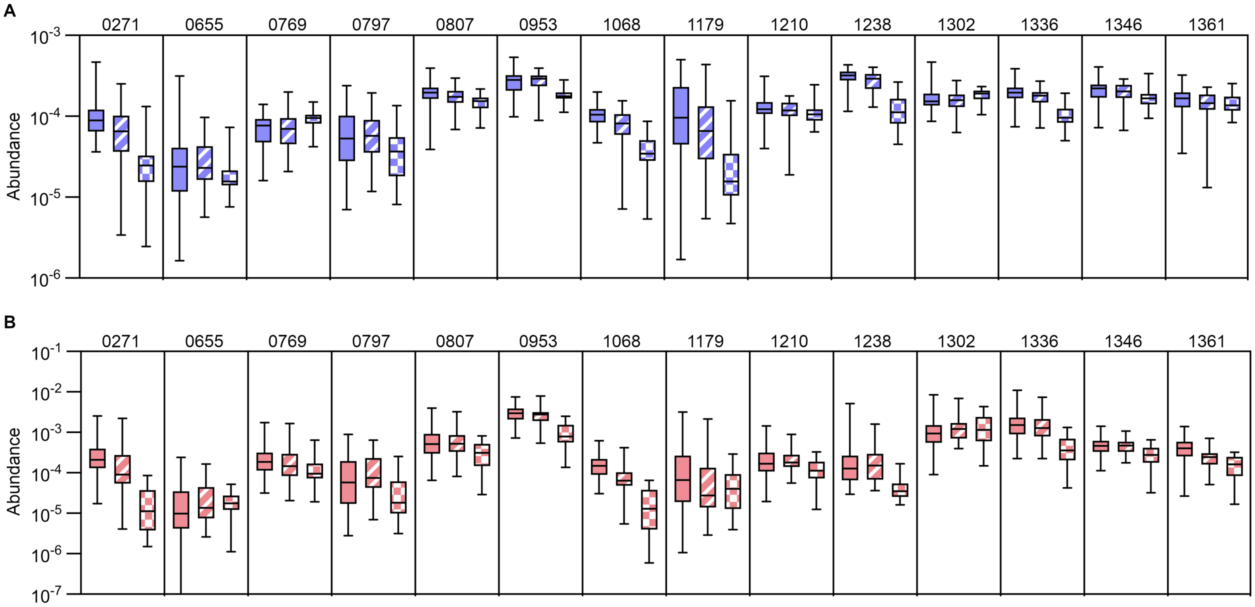


C

| Protein | Metagenome: SRF vs. DCM | Metagenome: SRF vs. MES | Metatranscriptome: SRF vs. DCM | Metatranscriptome: SRF vs. MES |
|---|---|---|---|---|
| SAR11_0271 | 0.0076 ** | <0.0001 **** | 0.0477 * | <0.0001 **** |
| SAR11_0655 | 0.9142 | 0.3109 | 0.5643 | 0.4087 |
| SAR11_0769 | 0.9817 | <0.0001 **** | 0.8130 | 0.0054 ** |
| SAR11_0797 | 0.9344 | 0.0570 | 0.6683 | 0.0184 * |
| SAR11_0807 | 0.4603 | <0.0001 **** | 0.9888 | 0.0054 ** |
| SAR11_0953 | 0.9817 | <0.0001 **** | 0.7816 | <0.0001 **** |
| SAR11_1068 | 0.0064 ** | <0.0001 **** | <0.0001 **** | <0.0001 **** |
| SAR11_1179 | 0.4462 | <0.0001 **** | 0.2969 | 0.0397 * |
| SAR11_1210 | 0.8706 | 0.0833 | 0.9755 | 0.0166 * |
| SAR11_1238 | 0.3175 | <0.0001 **** | 0.9888 | <0.0001 **** |
| SAR11_1302 | 0.9344 | 0.1595 | 0.4133 | 0.5430 |
| SAR11_1336 | 0.2181 | <0.0001 **** | 0.7816 | <0.0001 **** |
| SAR11_1346 | 0.9344 | 0.0042 ** | 0.9888 | 0.0031 ** |
| SAR11_1361 | 0.8706 | 0.3109 | <0.0001 **** | <0.0001 |

**Fig. S9. Comparison of SBP abundance across ocean layers.** (**A-B**) Box-and-whisker plots comparing abundance of each SBP in epipelagic/surface (SRF, solid color), deep chlorophyll maximum (DCM, stripes), and mesopelagic (MES, checkerboard) samples from the (**A**) OM-RGCv2+G metagenome and (**B**) OM-RGCv2+T metatranscriptome datasets. Abundance data was obtained as described in **Fig. S7** and **Fig. S8**. (**C**) *P* values for comparison of mean abundance in SRF samples with mean abundance in DCM and MES samples. Significance was evaluated using Welch's unequal variances *t*-test with the Holm-Šídák correction for multiple comparisons using a significance threshold of $P < 0.05$. Orange shading indicates higher abundance in the SRF sample, while green shading indicates higher abundance in the DCM or MES sample.

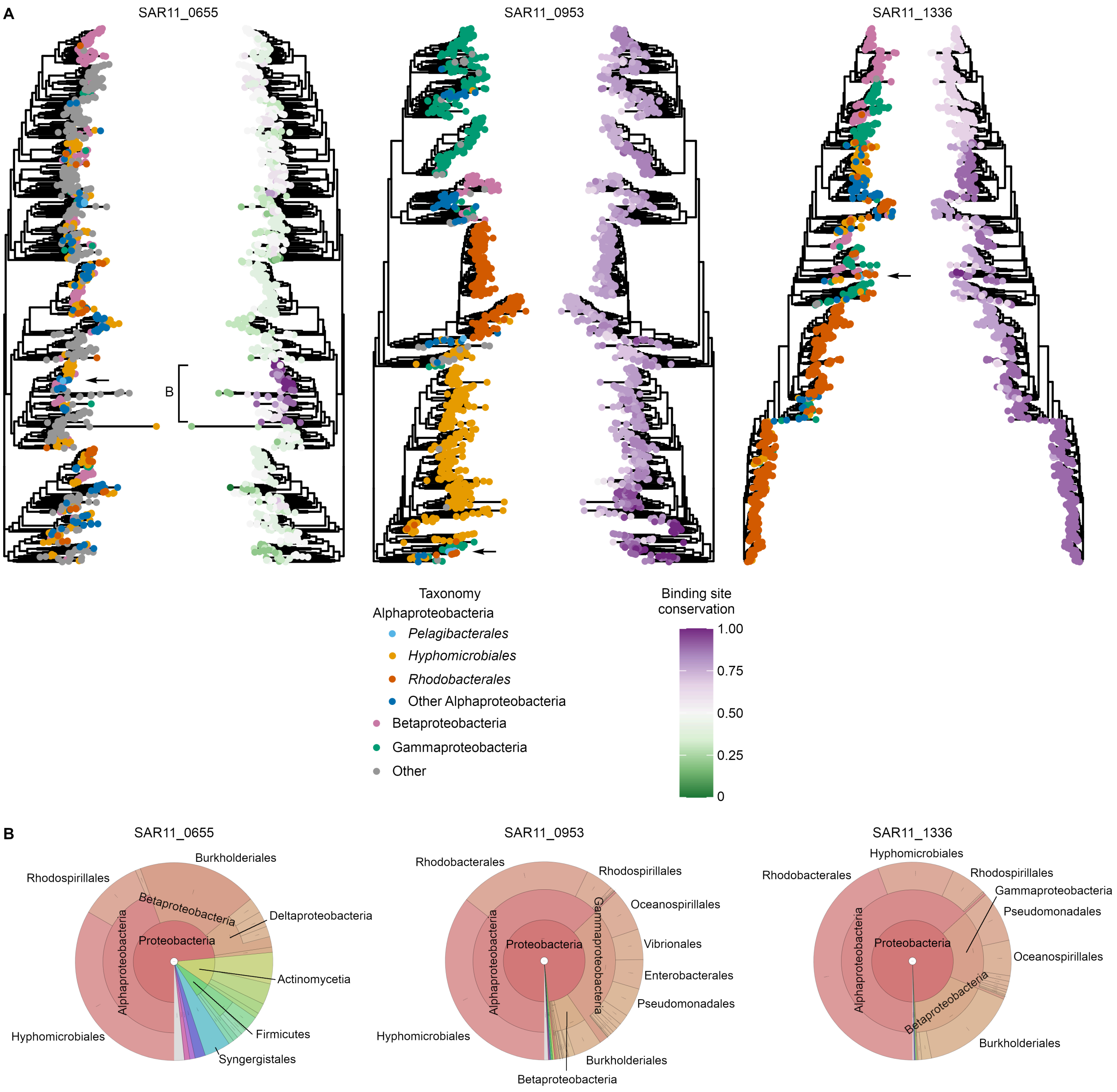


**Fig. S10. Phylogenetic analysis of selected SBPs from '*Ca.* P. ubique' HTCC1062.** (**A**) Maximum-likelihood phylogenies of 1000 homologs of SAR11_0655, SAR11_0953, and SAR11_1336 from the UniProtKB Reference Proteomes and Swiss-Prot databases. The positions of the SAR11 SBPs are indicated by an arrow. Nodes are colored by taxonomy and the fraction of binding site residues conserved relative to the corresponding protein in '*Ca*. P. ubique' HTCC1062. (**B**) Taxonomic distribution of sequences displayed in (A). In the case of SAR11_0655, only sequences belonging to the clade indicated in (A) were considered. Plots were generated using Krona[15].

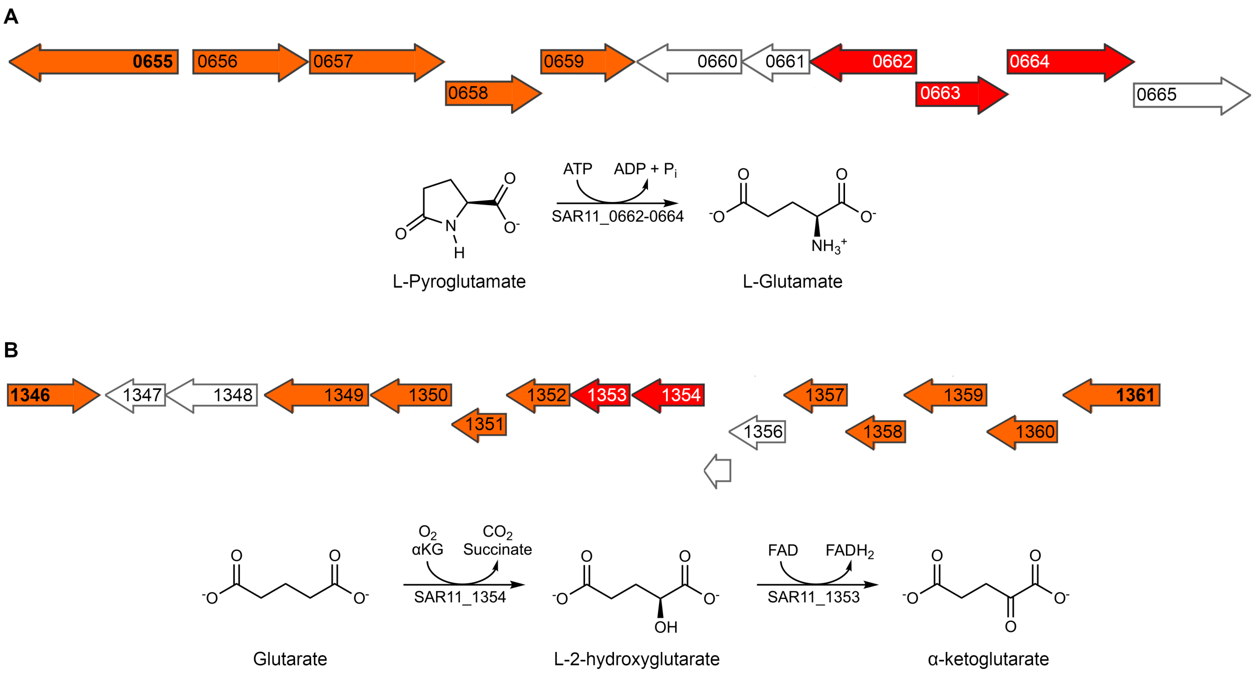


**Fig. S11. Genome context of SAR11_0655 and SAR11_1361 suggests additional metabolic capabilities of '*Ca*. P. ubique' HTCC1062.** ATP transporter genes (including SBP genes) are shown in orange, while genes putatively involved in metabolism of the transported substrates are shown in red. The genomic regions shown are bounded by non-coding regions of ≥75 bp. (**A**) Genome context of SAR11_0655 (9,248 bp). SAR11_0662–SAR11_0664 are homologous to the *pxpABC* genes from *E. coli* (sequence identity 30.0% overall). *pxpABC* encodes 5-oxoprolinase, which catalyzes ATP-dependent hydrolysis of L-pyroglutamate to L-glutamate[16]. (**B**) Genome context of SAR11_1346 and SAR11_1361 (14,635 bp). SAR11_1354 shows high sequence identity (41.5%) to *csiD* from *E. coli* encoding glutarate 2-hydroxylase, which converts glutarate to L-2-hydroxyglutarate and is involved in catabolism of L-lysine to α-ketoglutarate[17]. Although the operon in '*Ca*. P. ubique' is lacking the remainder of the the L-lysine catabolic pathway, it does contain a FAD-dependent oxidoreductase of unknown function (SAR11_1353), which may convert L-2-hydroxyglutarate to α-ketoglutarate by analogy with the L-lysine catabolic pathway. Abbreviations: αKG, α-ketoglutarate.

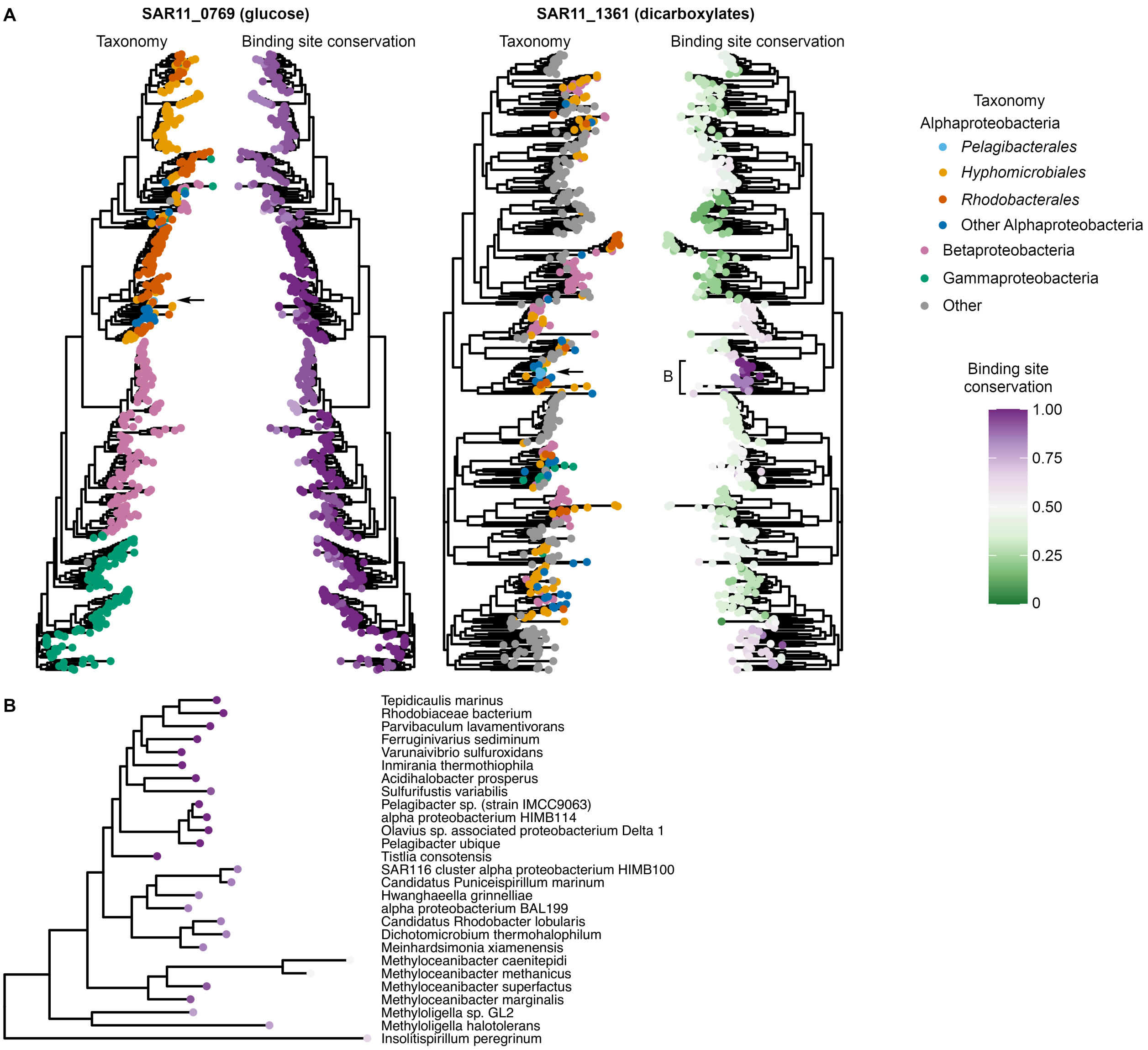


**Fig. S12. Contrasting phylogenetic distributions of SAR11_0769 and SAR11_1361.** (**A**) Maximum-likelihood phylogenies of 500 homologs of SAR11_0769 and SAR11_1361 from the UniProtKB Reference Proteomes and Swiss-Prot databases. The positions of SAR11_0769 and SAR11_1361 are indicated by an arrow. Nodes are colored by taxonomy and the fraction of binding site residues conserved relative to the corresponding protein in '*Ca*. P. ubique' HTCC1062. SAR11_0769 is widely distributed among bacteria, while SAR11_1361 appears to be limited mainly to SAR11 bacteria and a small range of other marine Alphaproteobacteria. (**B**) Expanded view of a clade of the SAR11_1361 phylogeny (indicated in **A**) showing protein sequences with a similar binding site to SAR11_1361 (suggesting a similar function).

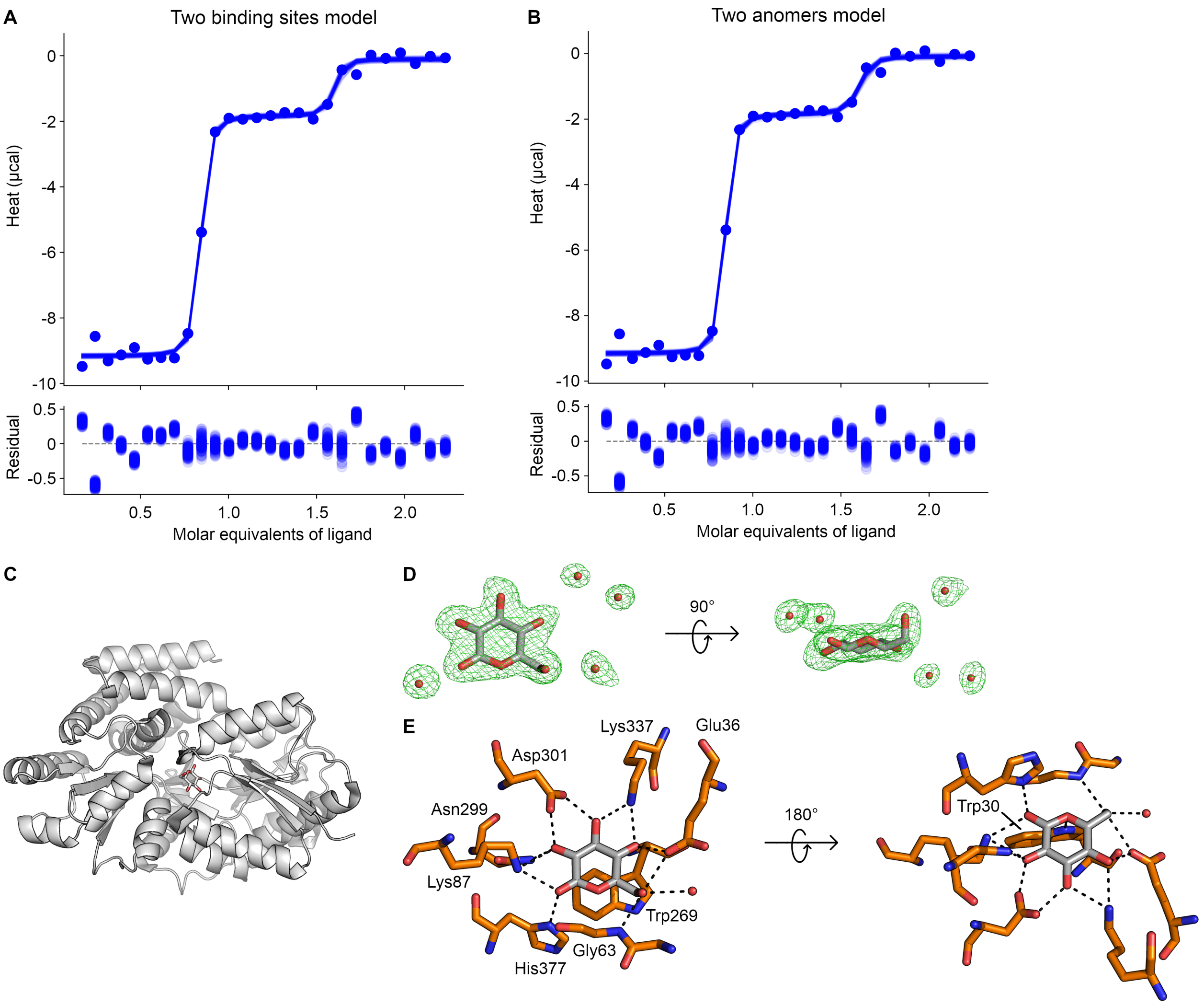


**Fig. S13. Binding mode of β-D-glucose to SAR11_0769. (A-B)** Representative ITC data for titration of SAR11_0769 with D-glucose, fitted to (**A**) the two-sets-of-sites binding model, or (**B**) a competitive binding model accounting for the two anomeric forms of D-glucose. Plots were generated using pytc[18]. (**C-E**) Crystal structure of SAR11_0769 complexed with β-D-glucose (1.86 Å). (**C**) Overall structure. (**D**) Electron density for the β-D-glucose molecule and neighbouring water molecules, shown by an m$F_o$ – d$F_c$ map contoured at +3σ. The density for the anomeric hydroxyl group is clearly resolved. (**E**) Binding mode of β-D-glucose.

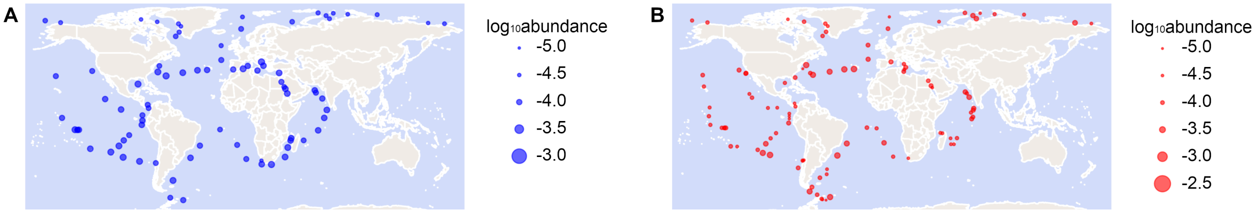


**Fig. S14. Broad geographical distribution of L-pyroglutamate-binding TRAP SBPs.** The *Tara* Oceans OM-RGCv2+G metagenome (**A**) and OM-RGCv2+T metatranscriptome (**B**) datasets were searched using Ocean Gene Atlas v2.0 with an e-value cut-off of $10^{-40}$ using the putative L-pyroglutamate-binding TRAP SBP from '*Ca*. P. ubique' HTCC9565 (NCBI accession WP_169035489.1) as the query sequence. Abundance at each location is expressed as the percentage of mapped reads and represented by point area on a linear scale.

**Table S1. Putative annotations of '*Ca*. P. ubique' SBPs prior to functional characterization.** Annotations were obtained from the NCBI Gene database, TransportDB 2.0 (a specialized database for annotation of transport proteins)[19], and UniProt. NCBI annotations are given at the time of commencing the project (old; January 2020) and after a more recent update (new; January 2022). The putative annotation in the final column represents our estimate of the consensus annotation. The closest homolog of each protein in the SwissProt and Protein Data Bank databases (January 2022) is also listed.

| Protein | New locus tag | UniProt ID | Annotations | | Closest homolog in SwissProt | | Closest homolog in PDB | | Putative annotation |
|---|---|---|---|---|---|---|---|---|---|
| SAR11_0271 | SAR11_RS01350 | Q4FNZ7 | NCBI (new) | ABC transporter substrate-binding protein | ID | Q7LYW7 | ID | 1EU8 | Sugar |
| | | | NCBI (old) | ABC transporter substrate-binding protein | Name | Trehalose/maltose-binding protein MalE | Name | Trehalose/maltose-binding protein MalE | |
| | | | TCDB | Sugar | Organism | *Thermococcus litoralis* | Organism | *Thermococcus litoralis* | |
| | | | UniProt | Putative ABC transporter solute-binding protein | E-value | 3e-14 | E-value | 8e-15 | |
| SAR11_0655 | SAR11_RS03320 | Q4FMW4 | NCBI (new) | ABC transporter substrate-binding protein | ID | Q8FWQ3 | ID | 3I45 | Branched-chain amino acids |
| | | | NCBI (old) | Leu/Ile/Val-binding protein | Name | Leu/Ile/Val-binding protein homolog 8, BRA0392 | Name | Twin-arginine translocation pathway signal, Rru_A0563 | |
| | | | TCDB | Leucine/valine | Organism | *Brucella suis* | Organism | *Rhodospirillum rubrum* | |
| | | | UniProt | Probable Leu/Ile/Val-binding protein *braC* | E-value | 4e-45 | E-value | 0.003 | |
| SAR11_0769 | SAR11_RS03875 | Q4FMK2 | NCBI (new) | ABC transporter substrate-binding protein | ID | O06875 | ID | 4R2B | Sugar |
| | | | NCBI (old) | Sugar ABC transporter substrate-binding protein | Name | Probable sugar-binding periplasmic protein BruAb2_0537 | Name | Extracellular solute-binding protein family 1 (glucose-bound) | |
| | | | TCDB | Sugar | Organism | *Brucella abortus* | Organism | *Brucella anthropi* | |
| | | | UniProt | Probable binding protein component of ABC sugar transporter | E-value | 2e-127 | E-value | 4e-127 | |
| SAR11_0797 | SAR11_RS04015 | Q4FMH4 | NCBI (new) | Glycine/betaine ABC transporter substrate-binding protein | No hits | | No hits | | Glycine betaine |
| | | | NCBI (old) | Hypothetical protein | | | | | |
| | | | TCDB | Glycine betaine | | | | | |
| | | | UniProt | OpuAC domain-containing protein *proX* | | | | | |
| SAR11_0807 | SAR11_RS04065 | Q4FMG4 | NCBI (new) | ABC transporter substrate-binding protein | ID | P40400 | ID | 3UIF | Taurine |
| | | | NCBI (old) | Taurine transport system periplasmic protein | Name | Putative aliphatic sulfonates-binding protein ssuA | Name | Sulfonate ABC transporter, periplasmic sulfonate-binding protein SsuA, Mfla_1563 | |
| | | | TCDB | Nitrate/sulfonate/taurine | Organism | *Bacillus subtilis* | Organism | *Methylobacillus flagellatus* | |

| | | | | | | | | | |
|---|---|---|---|---|---|---|---|---|---|
| | | | UniProt | Taurine transport system periplasmic protein *tauA* | E-value | 5e-08 | E-value | 0.002 | |
| SAR11_0953 | SAR11_RS04760 | Q4FM26 | NCBI (new) | Amino acid ABC transporter substrate-binding protein | ID | Q52812 | ID | 4Z9N | Amino acids |
| | | | NCBI (old) | ABC transporter | Name | General L-amino acid-binding periplasmic protein AapJ | Name | Amino acid ABC transporter, periplasmic amino acid-binding protein BOV_0736 (glutathione-bound) | |
| | | | TCDB | Amino acid (glutamine/ glutamate/aspartate?) | Organism | *Rhizobium leguminosarum* | Organism | *Brucella ovis* | |
| | | | UniProt | ABC transporter *yhdW* | E-value | 6e-130 | E-value | 7e-132 | |
| SAR11_1068 | SAR11_RS05345 | Q4FLR5 | NCBI (new) | Transporter substrate-binding domain-containing protein | ID | Q01269 | ID | 5HPQ[1] | Cyclohexadien-yl dehydratase |
| | | | NCBI (old) | Cyclohexadienyl dehydratase | Name | Cyclohexadienyl dehydratase *pheC* | Name | Cyclohexadienyl dehydratase *pheC* | |
| | | | TCDB | Amino acid (glutamine/ glutamate/aspartate?) | Organism | *Pseudomonas aeruginosa* | Organism | *Pseudomonas aeruginosa* | |
| | | | UniProt | Cyclohexadienyl dehydratase | E-value | 6e-28 | E-value | 9e-29 | |
| SAR11_1179 | SAR11_RS05925 | Q4FLF3 | NCBI (new) | Substrate-binding domain-containing protein | ID | Q55200 | No hits | | Phosphate |
| | | | NCBI (old) | Phosphate ABC transporter | Name | Protein SphX (phosphate-binding protein) | | | |
| | | | TCDB | Not found | Organism | *Synechocystis sp.* | | | |
| | | | UniProt | Phosphate ABC transporter *pstS* | E-value | 3e-24 | | | |
| SAR11_1210 | SAR11_RS06075 | Q4FLC2 | NCBI (new) | Transporter substrate-binding domain-containing protein | ID | P72298 | ID | 5OT8 | Octopine/ nopaline |
| | | | NCBI (old) | ABC transporter *occT* | Name | Octopine-binding periplasmic protein *occT* | Name | Nopaline-binding periplasmic protein *nocT* (octopine-bound) | |
| | | | TCDB | Amino acid (glutamine/ glutamate/aspartate?) | Organism | *Rhizobium meliloti* | Organism | *Agrobacterium fabrum* | |
| | | | UniProt | ABC transporter *occT* | E-value | 7e-54 | E-value | 1e-52 | |
| SAR11_1238 | SAR11_RS06215 | Q4FL96 | NCBI (new) | Fe(3+) ABC transporter substrate-binding protein | ID | P72827 | ID | 2PT1 | Iron(III) |
| | | | NCBI (old) | Iron-uptake ABC transport system periplasmic iron-binding protein | Name | Iron uptake protein A1 *futA1* | Name | Iron uptake protein A1 *futA1* | |
| | | | TCDB | 2-aminoethylphosphonate | Organism | *Synechocystis sp.* | Organism | *Synechocystis sp.* | |
| | | | UniProt | Probable iron-uptake ABC transport system periplasmic iron-binding protein *sfuC* | E-value | 6e-94 | E-value | 2e-94 | |
| SAR11_1302 | SAR11_RS06515 | Q4FL33 | NCBI (new) | ABC transporter substrate-binding protein | No hits | | ID | 4XZ6 | Glycine betaine[2] |
| | | | NCBI (old) | Substrate-binding region of ABC-type glycine betaine transport system *opuAC* | | | Name | Glycine betaine/proline ABC transporter, periplasmic substrate-binding protein SPO1548 (trimethylamine-*N*-oxide-bound) | |
| | | | TCDB | Glycine betaine | | | Organism | *Ruegeria pomeroyi* | |

| | | | | | | | | | |
|---|---|---|---|---|---|---|---|---|---|
| | | | UniProt | Substrate-binding region of ABC-type glycine betaine transport system *opuAC* | | | E-value | 3e-92 | |
| SAR11_1336 | SAR11_RS06690 | Q4FKZ8 | NCBI (new) | Extracellular solute-binding protein | No hits | | ID | 4I1D | Spermidine/putrescine |
| | | | NCBI (old) | Spermidine/putrescine-binding periplasmic protein *potD* | | | Name | ABC transporter substrate-binding protein (unknown specificity) | |
| | | | TCDB | Spermidine/putrescine | | | Organism | *Bradyrhizobium diazoefficiens* | |
| | | | UniProt | Spermidine/putrescine-binding periplasmic protein *potD* | | | E-value | 1e-32 | |
| SAR11_1346 | SAR11_RS06735 | Q4FKY9 | NCBI (new) | ABC transporter substrate-binding protein | ID | Q8YEE8 | ID | 4N0Q | Branched-chain amino acids |
| | | | NCBI (old) | Hypothetical protein | Name | Leu/Ile/Val-binding protein homolog 3 | Name | Leu/Ile/Val-binding protein homolog 3 (leucine-bound) | |
| | | | TCDB | Leucine/valine | Organism | *Brucella melitensis* | Organism | *Brucella melitensis* | |
| | | | UniProt | Peripla_BP_6 domain-containing protein *livJ* | E-value | 2e-21 | E-value | 2e-20 | |
| SAR11_1361 | SAR11_RS06810 | Q4FKX4 | NCBI (new) | Amino acid ABC transporter substrate-binding protein | ID | Q2YJA9 | ID | 7S6F | Branched-chain amino acids |
| | | | NCBI (old) | Leu/Ile/Val-binding protein precursor *livJ2* | Name | Leu/Ile/Val-binding protein homolog 5 | Name | Putative urea ABC transporter, urea binding protein *urtA1* (urea-bound) | |
| | | | TCDB | Leucine/valine | Organism | *Brucella abortus* | Organism | *Parasynechococcus marenigrum* | |
| | | | UniProt | Leu/Ile/Val-binding protein *livJ2* | E-value | 8e-66 | E-value | 2e-15 | |

[1]Excluding synthetic proteins and SAR11_1068 itself.
[2]SAR11_1302 was independently shown to bind trimethylamine-*N*-oxide while this work was in progress [20].

**Table S2. Detection of '*Ca.* P. ubique' HTCC1062 SBPs in the Sargasso Sea.** The table shows literature data from the metaproteomic analysis of Sowell *et al*. (2009)[21]. The number of LC/MS-MS spectra is shown for each protein that is homologous to one of the SBPs found in '*Ca.* P. ubique' HTCC1062. Although the abundance of each SBP is not directly comparable, because '*Ca.* P. ubique' HTCC1062 is not the dominant SAR11 strain in the Sargasso Sea, these data provide evidence that most of the SBPs are expressed to some extent in SAR11 bacteria under environmentally relevant conditions. n.d. not detected.

| **ID** | **HTCC1062 equivalent** | **Number of spectra** |
|---|---|---|
| PB7211_1190 | SAR11_1179 | 2949 |
| PB7211_1204 | SAR11_0953 | 1479 |
| PB7211_697 | SAR11_1336 | 497 |
| PB7211_687 | SAR11_1302 | 367 |
| PB7211_1324 | SAR11_1361 | 360 |
| PB7211_601 | SAR11_0807 | 184 |
| PB7211_130 | SAR11_1346 | 148 |
| YP_266190 | SAR11_0769 | 84 |
| YP_266078 | SAR11_0655 | 28 |
| PB7211_689 | SAR11_1210 | 26 |
| PB7211_754 | SAR11_0797 | 22 |
| PB7211_704 | SAR11_1068 | 11 |
| | SAR11_0271 | n.d. |
| | SAR11_1238 | n.d. |

**Table S3. (separate file)**
**Full list of ligands used for high-throughput screening of SBP function.**

**Table S4. List of hits from Biolog assays and in-house screen.** All ligands that gave a $\Delta T_M \geq 2$ °C are shown. $\Delta T_M$ values are derived from a single replicate experiment and are not directly comparable between wells due to differences in concentration. Some compounds are found in multiple Biolog microarray plates and are therefore listed multiple times.

| Protein | Well | $\Delta T_m$ (°C) | Compound |
|---|---|---|---|
| **SAR11_0655** | PM3-D3 | 13.89 | L-pyroglutamic acid |
| | PM2-H3 | 10.82 | L-pyroglutamic acid |
| | PM1-E1 | 8.67 | L-glutamine |
| | PM3-B1 | 7.71 | L-glutamine |
| | PM4-G5 | 4.56 | Glutathione |
| **SAR11_0769** | PM1-C9 | 15.17 | α-D-glucose |
| | PM3-E6 | 13.94 | Glucuronamide |
| | PM1-E10 | 7.48 | Maltotriose |
| | - | 7.18 | L-arabinose |
| | PM4-F11 | 6.43 | Cysteamine |
| | PM2-A6 | 6.32 | Dextrin |
| | PM4-D3 | 6.21 | Cysteamine-*S*-phosphate |
| | PM1-A11 | 6.19 | D-mannose |
| | PM1-B8 | 6.14 | D-xylose |
| | PM4-F4 | 5.69 | Tetrathionate |
| | PM2-C8 | 5.67 | 3-methylglucose |
| | PM4-C5 | 5.49 | 2-deoxy-D-glucose-6-phosphate |
| | PM1-F11 | 5.13 | D-cellobiose |
| | PM4-F3 | 5.04 | Thiosulfate |
| | PM2-D7 | 4.13 | Turanose |
| | PM2-A11 | 4.05 | Mannan |
| | PM1-A6 | 3.69 | D-galactose |
| | PM2-C12 | 3.66 | Palatinose |
| | PM1-E8 | 3.44 | β-methyl-D-glucoside |
| | PM1-H7 | 3.34 | Glucuronamide |
| | PM3-E10 | 3.28 | D-mannosamine |
| | PM2-A12 | 3.19 | Pectin |
| | PM3-E8 | 3.14 | D-glucosamine |
| | PM2-C4 | 2.61 | D-melezitose |
| | PM2-B11 | 2.57 | D-fucose |
| | PM2-B12 | 2.46 | 3-*O*-β-D-galactopyranosyl-D-arabinose |
| | PM1-A10 | 2.39 | D-trehalose |
| | PM2-B3 | 2.32 | β-D-allose |
| **SAR11_0797** | - | 7.47 | Choline-*O*-sulfate |
| | - | 6.71 | Phosphorylcholine |
| | - | 4.27 | Betaine |
| | PM4-E4 | 2.79 | Phosphorylcholine |
| | - | 2.34 | DMSP |
| | - | 2.07 | Thiamine pyrophosphate |

| | | | |
|---|---|---|---|
| **SAR11_0807** | PM4-H6 | 12.85 | Taurine |
| | PM4-H7 | 7.79 | Hypotaurine |
| | PM4-F4 | 5.64 | Tetrathionate |
| | PM2-A8 | 5.39 | Glycogen |
| | - | 3.47 | Choline-*O*-sulfate |
| | PM1-B7 | 2.13 | D,L-α-glycerol-phosphate |
| **SAR11_0953** | PM3-A12 | 12.45 | L-glutamic acid |
| | PM3-A9 | 9.68 | L-asparagine |
| | PM1-B12 | 8.39 | L-glutamic acid |
| | PM3-A10 | 8.09 | L-aspartic acid |
| | PM1-D1 | 6.70 | L-asparagine |
| | PM3-G12 | 6.59 | α-aminovaleric acid |
| | PM3-B3 | 6.49 | L-histidine |
| | PM3-D2 | 6.45 | N-phthaloyl-L-glutamic acid |
| | PM3-A11 | 5.46 | L-cysteine |
| | PM3-G7 | 5.03 | D,L-α-aminobutyric acid |
| | PM3-C11 | 5.02 | L-homoserine |
| | PM3-B5 | 4.92 | L-leucine |
| | PM3-B11 | 4.69 | L-threonine |
| | PM1-A7 | 4.63 | L-aspartic acid |
| | PM3-B10 | 4.22 | L-serine |
| | PM2-G6 | 3.19 | L-histidine |
| | PM3-C2 | 3.07 | L-valine |
| | PM1-G4 | 2.63 | L-threonine |
| | PM2-G7 | 2.62 | L-homoserine |
| | PM2-G10 | 2.49 | L-leucine |
| | PM3-A7 | 2.46 | L-alanine |
| | PM3-B7 | 2.38 | L-methionine |
| | PM4-E4 | 2.29 | Phosphorylcholine |
| | PM3-B1 | 2.29 | L-glutamine |
| | - | 2.23 | Proline betaine |
| **SAR11_1068** | N/A | N/A | N/A (none identified) |
| **SAR11_1210** | PM3-A8 | 8.41 | L-arginine |
| | PM3-B6 | 7.07 | L-lysine |
| | PM4-D4 | 5.65 | Phospho-L-arginine |
| | PM2-G4 | 4.72 | L-arginine |
| | PM2-G11 | 3.97 | L-lysine |
| | PM2-A11 | 3.56 | Mannan |
| | - | 2.30 | D-octopine |
| **SAR11_1302** | - | 18.65 | Trimethylamine-N-oxide |
| | - | 12.38 | Choline |
| **SAR11_1336** | - | 13.92 | Ectoine |
| | - | 13.42 | Betaine |
| | - | 13.25 | DMSP |
| | - | 13.10 | Trigonelline |
| | - | 11.54 | Proline betaine |

| | | | |
|---|---|---|---|
| | - | 11.54 | Pipecolate |
| | PM3-B9 | 8.88 | L-proline |
| | PM1-A8 | 6.96 | L-proline |
| | PM3-G8 | 5.32 | γ-aminobutyric acid |
| | PM3-C3 | 4.46 | D-alanine |
| | PM2-D10 | 3.97 | γ-aminobutyric acid |
| | PM1-A9 | 3.61 | D-alanine |
| | PM3-G11 | 3.57 | δ-aminovaleric acid |
| | PM3-G7 | 2.52 | DL-α-aminobutyric acid |
| | PM3-A7 | 2.24 | L-alanine |
| | PM3-G9 | 2.07 | ε-aminocaproic acid |
| **SAR11_1361** | PM4-F4 | 14.59 | Tetrathionate |
| | PM1-D6 | 13.50 | α-ketoglutaric acid |
| | PM1-G11 | 10.33 | D-malic acid |
| | PM1-A5 | 9.69 | Succinic acid |
| | PM2-E11 | 9.04 | Itaconic acid |
| | PM1-C3 | 8.18 | D,L-malic acid |
| | PM2-F12 | 8.08 | L-tartaric acid |
| | PM3-A12 | 7.60 | L-glutamic acid |
| | PM1-F5 | 7.11 | Fumaric acid |
| | PM1-G12 | 7.10 | L-malic acid |
| | PM3-F12 | 6.54 | Inosine |
| | PM1-F6 | 6.49 | Bromosuccinic acid |
| | PM1-G9 | 5.79 | Monomethyl succinate |
| | PM1-E6 | 5.53 | α-hydroxyglutaric acid-γ-lactone |
| | PM4-F3 | 5.01 | Thiosulfate |
| | PM2-F5 | 4.48 | Oxalomalic acid |
| | PM2-F9 | 4.01 | Sorbic acid |
| | PM2-F10 | 3.52 | Succinamic acid |
| | PM1-G2 | 3.27 | Tricarballylic acid |
| | PM3-A10 | 3.24 | L-aspartic acid |
| | PM3-C5 | 3.15 | D-aspartic acid |
| | PM3-G4 | 3.00 | Alloxan |
| | PM2-E4 | 2.70 | Citramalic acid |
| | PM1-E2 | 2.48 | *meso*-tartaric acid |
| | PM2-F11 | 2.23 | D-tartaric acid |

**Table S5. Binding parameters for SBP-ligand interactions derived from ITC.** The fraction of active protein ($n$), dissociation constant ($K_d$), and binding enthalpy ($\Delta H$) is given for each interaction. Data represent mean ± s.d. for the given number of technical replicates (#; separate titrations). For some titrations, only an upper limit on $K_d$ could be estimated. Fitted values of $n$, $K_d$ and $\Delta H$ together with fitting errors for individual titrations are given in **Data S1**. n.d. binding not detected.

| Protein | Ligand | # | *n* | $K_d$ (nM) | $\Delta H$ (kcal/mol) |
|---|---|---|---|---|---|
| **SAR11_0655** | Pyroglutamate | 3 | 0.33 ± 0.02 | < 5 | +4.5 ± 0.0 |
| | Glutamine | 1 | n.d. | n.d. | n.d. |
| **SAR11_0769** | Glucose[1] | 4 | | | |
| | High-affinity anomer | | 0.77 ± 0.07 | < 0.027 | -25.0 ± 0.3 |
| | Low-affinity anomer | | | 37 ± 20 | -17.4 ± 0.8 |
| | Galactose[2] | 3 | - | - | - |
| | Glucosamine[3] | 1 | 1.28 ± 0.03 | 238 ± 132 | -4.9 ± 0.3 |
| | Arabinose | 2 | 0.90 ± 0.02 | 1310 ± 90 | -11.2 ± 0.1 |
| | Maltotriose | 1 | n.d. | n.d. | n.d. |
| | Cellobiose | 1 | n.d. | n.d. | n.d. |
| **SAR11_0797** | Choline-*O*-sulfate | 2 | n.d. | n.d. | n.d. |
| | Phosphocholine | 2 | n.d. | n.d. | n.d. |
| | Glycine betaine | 1 | n.d. | n.d. | n.d. |
| **SAR11_0807** | Taurine | 3 | 0.98 ± 0.08 | 62 ± 12 | -12.9 ± 0.2 |
| **SAR11_0953** | Glutamate | 3 | | | |
| | Direct | | 0.37 ± 0.05 | < 5 | -13.1 ± 0.2 |
| | Competition (phenylalanine) | | 0.47 ± 0.02 | 0.55 ± 0.32 | -12.1 ± 0.4 |
| | Serine | 2 | 0.40 ± 0.02 | 6.4 ± 0.6 | -15.2 ± 0.8 |
| | Glycine | 2 | 0.39 ± 0.04 | 26 ± 5 | -17.1 ± 0.6 |
| | Lysine | 2 | 0.42 ± 0.03 | 195 ± 126 | -16.9 ± 0.4 |
| | Phenylalanine[4] | 6 | 0.41 ± 0.3 | 5530 ± 1750 | -5.0 ± 0.6 |
| **SAR11_1179** | Phosphate | 3 | 0.76 ± 0.03 | 133 ± 28 | -5.1 ± 0.6 |
| **SAR11_1210** | Arginine | | | | |
| | Direct | 3 | 0.67 ± 0.12 | < 5 | -4.9 ± 0.1 |
| | Competition (octopine) | 6 | 0.32 ± 0.13 | 0.058 ± 0.040 | -4.7 ± 0.3 |
| | Competition (SeArgT) | 2 | 0.63 ± 0.00 | 0.033 ± 0.014 | -4.4 ± 0.3 |
| | Octopine | 2 | 0.74 ± 0.01 | 232 ± 51 | +2.1 ± 0.1 |
| **SAR11_1336** | Glycine betaine | | | | |
| | Direct | 2 | 0.93 ± 0.02 | < 5 | -14.0 ± 0.1 |
| | Competition (glycine) | 2 | 0.95 ± 0.02 | 1.9 ± 0.1 | -13.3 ± 0.2 |
| | Glycine | 2 | 0.91 ± 0.16 | 5460 ± 60 | -6.1 ± 0.4 |

| | | | | | |
|---|---|---|---|---|---|
| **SAR11_1361** | α-Ketoglutarate | 2 | 0.82 ± 0.03 | 7.7 ± 1.4 | -15.7 ± 0.1 |
| | Oxaloacetate | 2 | 0.88 ± 0.04 | 9.6 ± 3.9 | -16.3 ± 0.1 |
| **SeArgT** | Arginine | | | | |
| | Direct | 3 | 0.88 ± 0.01 | 15 ± 7 | -11.3 ± 0.1 |
| | Competition (SAR11_1210) | 2 | 0.98 ± 0.01 | 10.0 ± 0.4 | -10.8 ± 0.1 |

[1]Reciprocal of mean lower bound of $K_a$ derived from Bayesian fitting.
[2]Binding was observed and the data were fitted to the two-sets-of-sites model, but the parameters of the model were highly correlated and unique values of $n$, $K_d$, and ΔH for each site could not be obtained.
[3]Fitted value ± standard error of fit for single replicate.
[4]Fitted value ± standard error of fit for single replicate with most stable baseline, out of 6 replicates.

**Table S6. Predicted and experimentally determined functions of the SBPs of '*Ca*. P. ubique' HTCC1062.** Further information about the functional predictions is given in **Table S1**.

| Protein | Predicted substrate/function | Confirmed substrate/function |
|---|---|---|
| SAR11_0271 | Sugars | Unknown |
| SAR11_0655 | Branched-chain amino acids | Pyroglutamate |
| SAR11_0769 | Sugars | Glucose |
| SAR11_0797 | Glycine betaine | Unknown |
| SAR11_0807 | Taurine | Taurine |
| SAR11_0953 | Amino acids | Amino acids |
| SAR11_1068 | Cyclohexadienyl dehydratase | Unknown |
| SAR11_1179 | Phosphate | Phosphate |
| SAR11_1210 | Octopine/nopaline | Arginine/lysine |
| SAR11_1238 | Iron(III) | Iron(III) |
| SAR11_1302 | Glycine betaine | Trimethylamine-*N*-oxide |
| SAR11_1336 | Spermidine/putrescine | Glycine betaine, DMSP and other osmolytes |
| SAR11_1346 | Branched-chain amino acids | Unknown |
| SAR11_1361 | Branched-chain amino acids | Dicarboxylates |

**Table S7. Literature measurements of ocean concentrations for selected nutrients.**

| Nutrient | Sample details | Concentration (nM) | Ref. |
|---|---|---|---|
| Amino acids | Surface Sargasso Sea, July (range, *n* = 14) | Glutamate **(<0.1–1.6**), Serine (**0.5–10.7**), Glutamine/histidine **(<0.1–1.6**), Glycine (**1.1–11.0**), Alanine (**0.5–5.7**), Leucine **(<0.1–1.7**), Total (**3.4–55.9**). | 22 |
| | Surface Atlantic Ocean, May and September-October, temperate provinces (mean ± s.d.; *n*) | Leucine (**0.23 ± 0.18**; 13), Methionine (**0.20 ± 0.11**; 13), Tyrosine (**0.18 ± 0.04**; 3). | 23 |
| | Surface Atlantic Ocean, May and September-October,, gyre provinces (mean ± s.d.; *n*) | Leucine (**0.21 ± 0.13**; 35), Methionine (**0.41 ± 0.18**; 41), Tyrosine (**0.16 ± 0.11**; 11). | 23 |
| | Surface Atlantic Ocean, May and September-October,, equatorial provinces (mean ± s.d.; *n*) | Leucine (**0.40 ± 0.34**; 13), Methionine (**0.69 ± 0.41**; 20), Tyrosine (**0.65 ± 0.57**; 8). | 23 |
| | Surface South China Sea, March to April (mean ± s.d.; range) | Arginine (**0.3 ± 0.8; 0.0–3.2**), Glutamate (**4.4 ± 2.9; 1.4–13**), Total (**95 ± 36; 52–202**) | 24 |
| | Mesopelagic South China Sea, March to April (mean ± s.d.; range) | Arginine (**0.5 ± 1.0; 0.0–2.9**), Glutamate (**5.5 ± 3.2; 0.8–11**), Total (**110 ± 41; 59–188**) | 24 |
| | Surface (5 m) North Atlantic Ocean near Iberian Peninsula, August (mean ± s.d.; range, *n* = 23) | Glutamate (**2.92 ± 2.06; 1.25–11.0**), Serine (**3.60 ± 2.29; 0.0–9.77**), Glycine (**4.08 ± 3.68; 0.04–14.5**), Arginine (**0.85 ± 0.73; 0–3.03**), Leucine (**0.77 ± 1.12; 0.12–5.6**), Lysine (**0.56 ± 0.76; 0.0–3.86**). (selected amino acids only) | 25 |
| | Baltic Sea, May, 1–75 m depth (mean ± s.d., *n* = 8) | Glutamate (**3.6 ± 1.0**), Serine (**18 ± 7**), Glycine (**13 ± 6**), Leucine (**1.2 ± 0.5**), Arginine (**3.8 ± 2.5**), Lysine (**5.8 ± 3.1**) | 26 |
| Taurine | Gulf of Alaska and North Atlantic, August | Approximate range of 1–10 in surface waters and 0.1–1 in mesopelagic waters | 27 |
| | Surface (5 m) North Atlantic Ocean near Iberian Peninsula, August (mean ± s.d.; range, *n* = 23) | **3.06 ± 3.15; 0.63–15.66** | 25 |
| DMSP | Surface Subarctic Pacific Ocean, July-August, coastal and oceanic samples (mean ± s.d.; range, *n* = 7) | **3.5 ± 2.7; 1.3–8.3** | 28 |
| | Surface Atlantic Ocean (40°S to 50°N), March-April (mean, range, *n*) | Area 1 (**3.66, 0.6–12.4**, 11), Area 2 (**1.38, 0.5–5.1**, 23), Area 3 (**1.03, 0.2–4.2**, 15), Area 4 (**1.6, 0.3–8.8**, 16), Area 5 (**11.4, 1.2–22.7**, 16) | 29 |

| | | | |
|---|---|---|---|
| Sugars | Surface Atlantic Ocean, May and September-October, temperate provinces (mean ± s.d.; *n*) | Glucose (**0.92 ± 0.62**; 8), *N*-Acetylglucosamine (**0.68 ± 0.21**; 3), Glucosamine (**0.39 ± 0.04**; 3). | [23] |
| | Surface Atlantic Ocean, May and September-October, gyre provinces (mean ± s.d.; *n*) | Glucose (**0.84 ± 0.47**; 11), *N*-Acetylglucosamine (**1.0 ± 0.54**; 7). | [23] |
| | Surface Atlantic Ocean, May and September-October, equatorial provinces (mean ± s.d.; *n*) | Glucose (**1.73 ± 0.67**; 4), *N*-Acetylglucosamine (**1.4 ± 0.53**; 4). | [23] |
| Phosphate | Surface North Atlantic subtropical gyre, bioavailable phosphate (mean ± s.d.; *n*) | September 2003 (**3.3 ± 1.2**; 3), May 2004 (**1.8 ± 1.1**; 4), October 2005 (**2.4 ± 1.3**; 15), September 2006 (**1.2 ± 0.3**; 4) | [30] |
| | Surface Sargasso Sea, March (mean ± s.d.?) | **0.48 ± 0.27** | [31] |
| | Surface North Pacific near Hawaii, November (mean ± s.d.?) | **13 ± 2** | [31] |

**Table S8. (separate file)**
**Sequences of oligonucleotides and synthetic genes used in this study.**

**Table S9. X-ray data collection and refinement statistics.**

| **Structure** | SAR11_0769/D-glucose | SAR11_1210/L-arginine |
|---|---|---|
| PDB code | 8HQQ | 8HQR |
| **Data collection** | | |
| Space group | $P3_221$ | $P2_12_12_1$ |
| Cell dimensions | | |
| *a*, *b*, *c* (Å) | 44.9, 44.9, 330.9 | 57.1, 84.7, 106.1 |
| *α*, *β*, *γ* (°) | 90.0, 90.0, 120.0 | 90.0, 90.0, 90.0 |
| Resolution (Å) | 47.02 – 1.86 (1.98 – 1.86) | 47.36 – 1.32 (1.40 – 1.32) |
| $R_{merge}$ (%) | 23.9 (52.1) | 12.3 (141.1) |
| $CC_{1/2}$ (%) | 98.6 (91.3) | 99.6 (70.9) |
| $I$ / σ($I$) | 6.28 (1.08) | 10.44 (1.16) |
| Completeness (%) | 99.0 (94.4) | 99.4 (99.1) |
| Redundancy | 12.0 (9.5) | 12.2 (11.6) |
| **Refinement** | | |
| Resolution (Å) | 38.40 – 1.86 | 47.4 – 1.32 |
| No. unique reflections | 33089 | 114968 |
| $R_{work}$ / $R_{free}$ (%) | 20.4 / 23.7 | 14.9 / 19.0 |
| Number of atoms (chain A / chain B) | | |
| Protein | 2960 | 1996 / 2005 |
| Ligand | 12 | 12 / 12 |
| Water | 99 | 578 |
| *B*-factors ($Å^2$) (chain A / chain B) | | |
| Protein | 34.9 | 23.4 / 22.6 |
| Ligand | 26.7 | 12.7 / 12.5 |
| Water | 33.4 | 34.0 |
| Root mean square deviations | | |
| Bond lengths (Å) | 0.007 | 0.009 |
| Bond angles (°) | 0.932 | 1.525 |

**Data S1. (separate file)**
**Data from individual ITC experiments.**

**Data S2. (separate file)**

**Binding affinities of previously reported SBPs.** In cases where multiple ligands were tested for binding, or multiple methods were used to measure binding affinity, the smallest dissociation constant (highest affinity) is given. Abbreviations: CD, circular dichroism; DSF, differential scanning fluorimetry; ED, equilibrium dialysis; IF, intrinsic tryptophan fluorescence; ITC, isothermal titration calorimetry; RA, radioassay; SPR, surface plasmon resonance.
